# Human hematopoietic development during the first year of life and its remodeling by neonatal Bacillus Calmette-Guérin vaccination

**DOI:** 10.64898/2026.11.25.768768

**Authors:** Lucinda Paddock, Leonie S. Helder, Chenyang Jiang, Collins K. Boahen, Nina Marie Birk Wischmann, Victoria Lawless, Alexia Martínez de Paz, Thomas Nørrelykke Nissen, Andrew W. Daman, Peter Aaby, Rachel E. Niec, Christine Stabell-Benn, Dana Pe’er, Mihai G. Netea, Steven Z. Josefowicz

**Author notes:** These authors contributed equally to this work (Lucinda Paddock, Leonie S. Helder, Chenyang Jiang).

## Abstract

Hematopoiesis and immune development are well defined in adults but remain poorly understood in early life, despite the neonatal period being critical for immune maturation. We map hematopoietic and immune programs across the first year of human life using single-nucleus epigenomic and transcriptomic profiling of circulating hematopoietic stem and progenitor cells (cHSPCs), including Bacillus Calmette–Guérin (BCG) vaccinated infants to illustrate a controlled immune challenge. BCG-naive infants displayed heightened myeloid activation through 3 months of age, followed by an interferon-enriched HSPC program at 13 months. BCG vaccination induced earlier interferon responses in HSPCs and dampened inflammatory programs in myeloid cells. These findings define the landscape of early-life human hematopoiesis and provide mechanistic insight into the heterologous protection associated with neonatal BCG vaccination.

---

The development of human immune cells from hematopoietic progenitors has been extensively profiled in adults and in the fetus (*1–5*), yet remains poorly understood in infants, despite the neonatal period being critical for immune maturation (*6–8*). During the first year of life, infants transition from a nearly sterile intrauterine environment to continuous microbial exposure, triggering considerable immune remodeling (*9, 10*). Their limited immune experience makes neonates highly vulnerable to pathogens that are much less virulent in adults, including microorganisms that can cause sepsis and respiratory infections, the leading contributors to infant mortality worldwide (*11*). Conversely, other infections with a poor prognosis in some adults, such as COVID-19, are most often mild in children (*12*). These fundamental differences between neonatal and adult immunity underscore the importance of the extensive hematopoietic and immune maturation that occurs during infancy, marked by coordinated shifts in immune-cell abundance and function (*13*). However, in-depth studies remain limited due to restricted access to infant biospecimens and the difficulty of conducting a prospective controlled study of immune response in newborns. As a result, developmental changes in human hematopoietic stem and progenitor cells (HSPCs) during the first year of life remain largely uncharacterized, including how they respond to environmental cues and how these influence mature immune cell phenotypes.

In the context of environmental and microbial exposures, standard immunizations provide a controlled and observable challenge to probe neonatal immune function (*14–16*). Bacillus Calmette–Guérin (BCG), consisting of live attenuated *Mycobacterium bovis*, is among the earliest and most widely administered vaccines, with 156 countries recommending BCG vaccination at birth in their vaccine schedule as of 2022 (*17*). BCG is typically delivered within days of birth to prevent severe tuberculosis and has long been associated with reduced mortality from heterologous infections (*18–24*), yet the mechanisms underlying these broader protective benefits in infants remain unclear.

Mouse studies demonstrate that BCG can reprogram hematopoiesis, inducing epigenetic remodeling of HSPCs that biases differentiation toward myeloid lineages with heightened antimicrobial capacity—a hallmark of trained immunity (*25*). Adult human studies reveal similar months-long transcriptional and epigenetic changes in HSPCs and their myeloid progeny following BCG vaccination (*26, 27*). Further, BCG immunotherapy for bladder cancer reprograms HSPC with a durable and transplantable capacity to generate myeloid cells with enhanced anti-tumor response (*28*). However, nearly all studies of BCG-induced trained immunity in humans have been conducted in adults, even though BCG is predominantly administered to neonates. How BCG influences early-life hematopoiesis and immune maturation remains unknown.

To address these questions, we applied Progenitor Input Enrichment combined single-nucleus RNA and ATAC (snRNA/ATAC) sequencing (PIE-Seq) to peripheral blood samples from children who participated in the Danish Calmette trial (*29*), a randomized clinical trial investigating the effects of BCG vaccination on childhood morbidity. This approach enables side-by-side profiling of circulating CD34^+^ progenitors and peripheral immune cells from small-volume infant blood samples (*28, 30*). From BCG-naive controls, we compiled a thorough timeline of progenitor and immune cell compositional changes from 4 days to 13 months of age, highlighting a shift from an early inflammatory bias to a progressive maturation of adaptive immunity accompanied by transcriptional and epigenetic changes in HSPC and myeloid compartments. These include substantial alterations in progenitor subset composition and transcriptional programs, with a gradual decline of a program of myeloid activation characterized by high-level NF-κB activity and a cumulative increase in an interferon gamma (IFN-γ) signature. We demonstrate that BCG vaccination induces earlier acquisition of the IFN-γ-dependent program, while curtailing early NF-κB-associated inflammatory programs in both HSPCs and myeloid cells. Together, these findings suggest a mechanistic model in which neonatal BCG vaccination reshapes infant hematopoiesis to promote a shift toward IFN-γ-driven immunity and away from myeloid NF-κB activation programs, providing a mechanistic model for heterologous protection associated with BCG.

## Distinct compositional profiles of human cord blood, infant PBMC, and adult PBMC

To examine the development of the immune system and hematopoiesis over the first year of human life, we analyzed cryopreserved PBMC samples from the Danish Calmette Study: a randomized clinical trial conducted from October 2012 to November 2015 investigating the effect of neonatal BCG vaccination on childhood morbidity in a cohort of 4,262 newborns (*29*). Infants were randomized within 7 days of birth to receive either BCG vaccination (Danish strain 1331) or no intervention (control group). Blood samples were collected from a subset of children (n=304) born in Hvidovre Hospital for immunological evaluation at 4 days (D4), 3 months (M3), and 13 months (M13) post-randomization. We employed PIE-Seq on matched samples from a subset of infants (n=12 BCG-vaccinated infants, n=10 BCG-naive or “control” infants), and independent adult (AD, n=7) and cord blood (CB, n=4) samples were included as reference cohorts. PIE-Seq enriches for rare circulating CD34^+^ HSPC (cHSPC) in peripheral blood before combined single-nucleus RNA and ATAC sequencing. A growing body of work has demonstrated that cHSPC capture the full heterogeneity and composition of bone marrow HSPC, and reflect disease, inflammation, and aging induced changes in hematopoiesis and immune progenitors (*28, 30–33*). PIE-Seq allows for side-by-side analysis of HSPC and mature cell gene expression and chromatin accessibility at single-nucleus resolution from a simple blood draw or cryopreserved PBMC, providing a unique opportunity to study cellular and molecular programs of neonatal hematopoiesis and bypassing ethical challenges and limitations in accessibility of neonatal bone marrow biopsies. These studies were complemented by functional analysis of whole-blood stimulation and serum protein data from the larger cohort of children in the Calmette trial (*34*). This allowed for integrated analysis of transcriptional and epigenetic programs with functional correlates of progenitor and immune phenotypes (Fig. 1A).

**Fig. 1.**
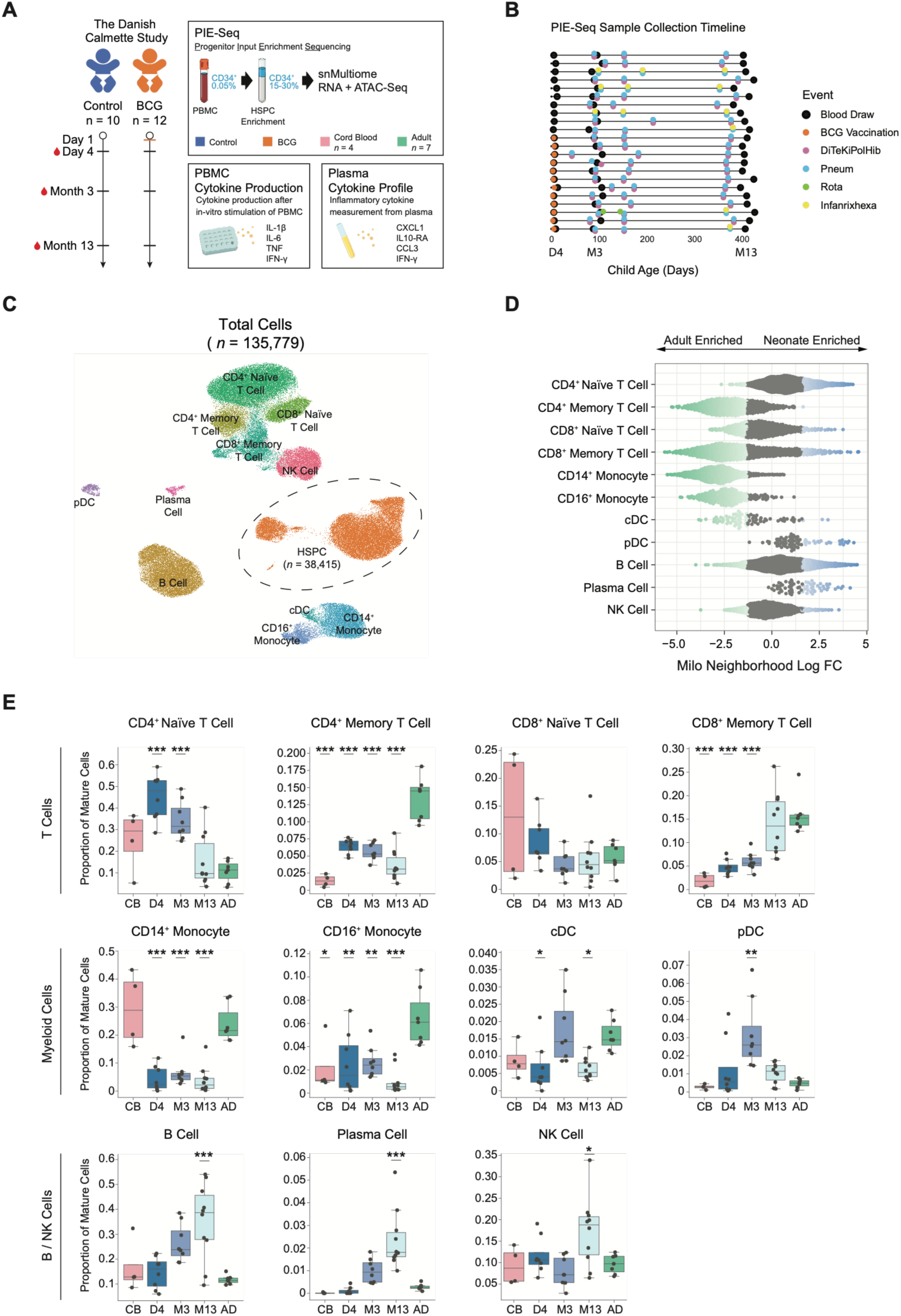
Profiling of infant immune maturation over the first year of life using PIE-Seq. (**A**) Study design: Samples from BCG-vaccinated and control infants were collected at 4 days, 3 months, and 13 months of age for profiling by PBMC paired single-nucleus ATAC- and RNA-seq with progenitor input enrichment (PIE-seq), PBMC cytokine stimulation assay, and serum inflammation panel. (**B**) Timeline of sample collection and administration of other childhood vaccines for PIE-seq samples. (**C**) UMAP visualization of mrVI *u* latent trained on snRNA data from all PIE-seq samples, labeled by cell type annotation with HSPC highlighted. (**D**) LogFC of Milo neighborhoods by cell type annotation, comparing all control infant to adult samples. Positive LogFC indicates local enrichment of infant cells. (**E**) Box plots of cell type proportion out of total mature cells for all control samples, colored by timepoint. Relative abundances were compared across timepoints by one-way ANOVA, with pairwise differences from the adult group determined by Tukey’s HSD test (adjusted P values shown). p > 0.05 = ns, ∗p ≤ 0.05, ∗∗p ≤ 0.01, ∗∗∗p ≤ 0.001.

Previous studies of the immune system in human infants have reported extensive heterogeneity between samples, with gestational age, chronological age, birth weight, and vaccine status as major sources of variation (*13, 35, 36*). Samples selected for PIE-Seq were representative of the overall cohort, with comparable birth weight and gestational age distribution between control and BCG-vaccinated infants (Fig. S1A,B), and were collected along a consistent timeline with minor variations in the administration of childhood vaccines (Fig. 1B, Table S1). Children in the BCG group were vaccinated within the first 7 days of life, and all children in the cohort were recommended the Danish standard childhood vaccination schedule.

Age was a dominant source of variation in both RNA and ATAC embeddings. Without integration, cells’ nearest neighbors in RNA embedding were predominantly drawn from the same age group, indicating that age-related variation outweighed cell-type identity at this level of representation (Fig. S1C,D). To enable consistent cell-type annotations while preserving biologically meaningful age effects, we applied MrVI to the snRNA data (*37*). MrVI models gene expression as a combination of cell-state–driven signals (u latent) and sample-associated variation (z latent), allowing these components to be disentangled. In the resulting cell-state (U) representation, age-driven neighborhood structure was substantially reduced (Fig. S1D), facilitating robust annotation across developmental stages. Using a classifier trained on adult reference data, we annotated 135,779 cells including 38,415 cHSPC (Fig. 1C). We found these cell type annotations to be robust to changes in MrVI model parameters, and concur with expression of well-known HSPC markers *MEIS1*, *CD34*, and *KIT* alongside canonical PBMC marker genes (Fig. S1E,F).

Initially we focused analysis on the non-BCG vaccinated control group, in order to understand dynamics in immune cells and HSPC across the neonatal period and adulthood in the absence of an introduced challenge. Given the observed age group effects, we applied the differential abundance method Milo to quantify changes in the composition of mature immune cells between samples from infants and adults (*38*). We found neighborhoods with significant enrichment of adult or infant cells across all annotated cell types, indicating broad changes in the makeup of peripheral blood immune cells over the first year of life (Fig. 1D, Fig. S1G). Visualizing cell type proportions out of total mature cells for each sample across all time points (Fig. 1E) revealed consistent and significant shifts over the first year of life, in addition to differences between cord blood, infant, and adult samples. The T-cell compartment (*THEMIS*^+^, *IL7R*^+^, and *CD3E*^+^) followed a broad trend of increasing antigenic experience, characterized by a decline in naive T-cells (*CD3E*^+^, *SELL*^high^ and *LEF1*^high^) and a corresponding rise in memory T-cells (*CD3E*^+^, *SELL*^low^ and *LEF1*^low^). By 13 months of age, infant CD8^+^ memory T-cells (*CD3E*^+^, *CD8A*^+^, and *SELL*^low^) were comparable to adult levels, while CD4^+^ memory T-cell proportions (*CD3E*^+^, *CD4*^+^, and *SELL*^low^) remained significantly lower, in line with the predominance of viral infections in infants. Within myeloid cells, both CD14^+^ (*LYZ*^+^ and *CD14*^+^) and CD16^+^ (*LYZ*^+^ and *FCGR3A*^+^) monocytes were consistently less abundant in infant blood than in adults, consistent with an established prominence of lymphopoiesis and reduced myelopoiesis in early life (*13, 39*). Interestingly, dendritic cell proportions displayed transient increases at 3 months of age for both conventional dendritic cells (cDC, *LYZ*^+^ and *HLA-DRA*^high^) and plasmacytoid dendritic cells (pDC, *CLEC4C*^+^ and *LILRA4*^+^). Future studies should assess if these DC trends might be associated with responses to specific developmental exposures that occur around 3 months. Finally, B cell abundance rose consistently during the first year of life, with both B cell (*EBF1*^+^ and *MS4A1*^+^) and plasma cell (*XBP1*^+^ and *PRDM1*^+^) numbers significantly elevated at 13 months compared to adults, similar to trends reported in previous studies of early-life immunity (*13, 40*).

Collectively, these findings illustrate that the first year of life is a period of significant changes in the human immune system. Immune cell proportions in infants not only differ from adults, but also display consistent developmental trajectories from a cord blood baseline to 13 months of age, underscoring the transition from a naive immune state to adult-like immunity. While these developmental trends at the level of the major immune cell populations are well-known (*13, 39*), this snRNA/ATAC-Seq PBMC dataset provides a unique cell atlas resource of circulating immune cell epigenetic and transcriptional programs from a cord-blood baseline through adulthood. Furthermore, with enrichment and analysis of HSPC, we can also interrogate the hematopoietic alterations that may underlie phenotypes observed in mature immune cells.

## Decreased plasticity of circulating HSPCs with age

We next characterized the cHSPC compartment in finer detail by re-embedding the snRNA layer of all annotated cHSPCs and applying finer cell filtering parameters to enable trajectory analysis. This snRNA/ATAC dataset of 36,417 CD34^+^ cHSPCs enabled us to perform in-depth analysis of hematopoiesis in human infants across the first year of life (Fig. 2A). To quantify changes in cHSPC subtype proportions, cHSPC clusters were annotated according to expression of a manually curated set of known HSPC lineage markers from our group’s previous work (*30*) (Fig. 2B). We found the cHSPC compartment in this dataset to be comprised of true hematopoietic stem cells/multipotent progenitors (HSC/MPP, hereafter referred to as HSC, *HLF^+^*and *CRHBP^+^*), lymphoid-primed multipotent progenitors (LMPP, *DNTT^+^* and *IRF8^+^*), granulocyte-monocyte progenitors (GMP, *MPO^+^* and *CEBPA^+^*), erythroid progenitors (ERP, *GATA1^+^* and *EPOR^+^*), basophil-eosinophil-mast cell progenitors (BEM, *HDC^+^* and *PRG2^+^*), and B-cell progenitors (BP, *EBF1^+^* and *CD79A^+^*). We further validated our annotations by calculating chromVAR motif accessibility of key hematopoietic transcription factors within HSPC subtypes (*41*), observing clear increases in motif accessibility for TCF family motifs along lymphoid fates, SPI1, SPIB and CEBPB family motifs along myeloid fates, and MECOM and GATA family motifs along erythroid fates (Fig. 2C).

**Fig. 2.**
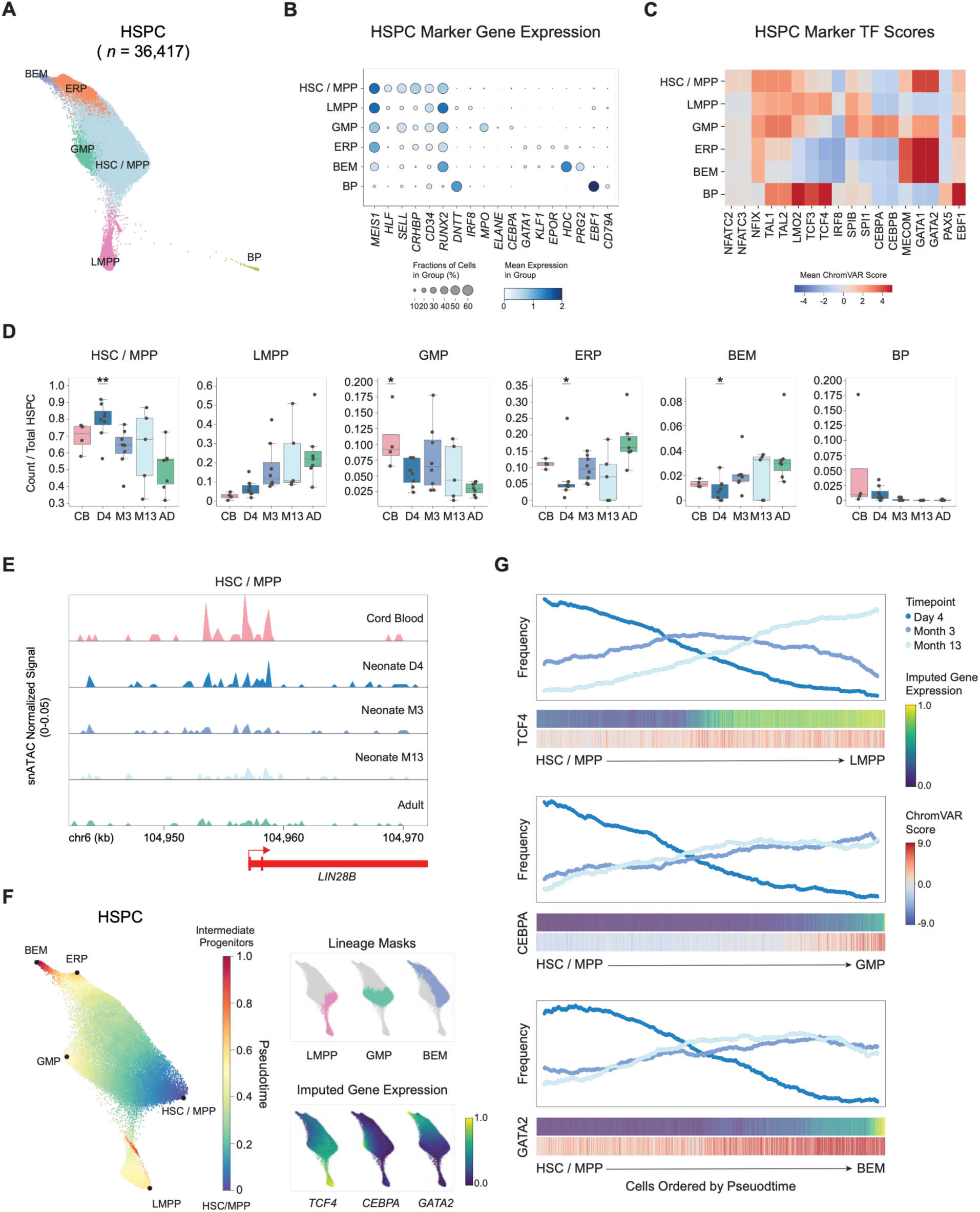
HSC plasticity in human infants decreases with age. (**A**) Force-directed layout (FDL) of snRNA data from all HSPC, colored by HSPC subtype cluster annotations. (**B**) Dot plot of marker gene expression (columns) by HSPC subtype annotation (rows). Dot size represents the number of cells per group expressing each gene, and dot color represents log-normalized expression level. (**C**) Heatmap of average ChromVAR score for marker transcription factor motifs (columns) across HSPC subtype annotations (rows). (**D**) Box plots of HSPC subtype proportion out of total HSPC for all control samples, colored by timepoint. Relative abundances were compared across timepoints by one-way ANOVA, with pairwise differences from the adult group determined by Tukey’s HSD test (adjusted P values shown). p > 0.05 = ns, ∗p ≤ 0.05, ∗∗p ≤ 0.01. (**E**) Pseudo-bulk ATAC-seq accessibility tracks of the *LIN28B* locus in HSC/MPP cells across conditions and timepoints. (**F**) (Left) Force-directed layout (FDL) of HSPC scRNA data excluding BP, colored by Palantir predicted pseudotime. Starting point (HSC / MPP) and end point (LMPP, GMP, ERP, BEM) cells are labeled. (Right) The same FDL colored by lineage branch masks generated from Palantir cell fate probabilities, and MAGIC-imputed expression of corresponding lineage marker genes. (**G**) Frequency of cells across pseudotime within each lineage branch mask, in bins of n = 1000 cells. Below each frequency plot is a heatmap of MAGIC imputed gene expression and chromVAR score for corresponding lineage markers over pseudotime-ordered cells.

Having established cHSPC subcluster annotations, we then investigated how these populations shift with age. Looking at proportions of subcluster annotations out of total cHSPCs, we observed striking changes in the HSC compartment, which at the Day 4 time point makes up an average of 79.1% of total cHSPCs before dropping to 50.8% in adults (Fig. 2D). Conversely, the more differentiated LMPP, ERP, and BEM populations display an increasing trend in abundance over the first year of life, especially between the Day 4 and Month 3 timepoints. Of all HSPC subtypes in our data, we found the BP cluster to be composed almost entirely of cord blood and infant cells (Fig. S2A), with most infant BP cells deriving from the Day 4 timepoint (Fig. 2D). The abundance of these cells in cord blood and Day 4 samples as compared to other timepoints suggests that this population is most relevant in the fetal or neonatal period and quickly decreases in circulation after birth.

We therefore characterized marker genes and motifs for the BP population in finer detail. Differential gene expression results comparing BPs to the rest of the cHSPC compartment indicated high expression of Pro-B cell transcriptional markers such as *DNTT*, *VPREB1*, *PAX5*, and *RAG1/2* (*42*) (Fig. S2B, Table S2). BPs also lacked expression of *CD19* and *MS4A1* (*CD20*), which identify more differentiated B-cell populations (Fig. S2B). Furthermore, BP marker peaks from the snATAC data layer were highly enriched for EBF1 and LYL1 motifs, which are both key factors in B-lymphopoiesis (*43*) (Fig. S2C, Table S2). Given the abundance of BPs in cord blood and infant samples relative to adults, we wondered whether they might be related to innate-like B-1 cells—a rare B-cell population that can differentiate from the embryonic yolk sac or fetal liver HSC, but not adult BM HSC (*44*). Experiments in mice have associated the ability of HSC to produce B-1 cells with expression of *Lin28b*, *Igf2bp1-3* and *Hmga2* (*45*). Indeed, we saw higher expression of their human orthologs in HSCs at early infant timepoints compared to adults in our data (Fig. S2D). Furthermore, we observed a marked decrease in *LIN28B* promoter accessibility in HSC from cord blood to later timepoints, in line with descriptions of murine *Lin28b* as a central regulator of the transition from fetal to postnatal hematopoiesis (*46*) (Fig. 2E). Therefore, we propose that the BP population likely has the potential to differentiate into B-1 cells. Our result suggests an early wave of distinct innate-like B cell progenitors, programmed at the level of early life HSC bias, that resolves before three months of age.

To better understand changes in HSPC differentiation in infants, we inferred cell fate trajectories using Palantir (*3*), which orders cells along a high-resolution pseudotime and estimates their relative bias toward distinct terminal lineages. We chose to exclude BP from this analysis given their separation from other cHSPC subclusters. Palantir ordered cells along expected differentiation trajectories, with HSC cells diverging towards LMPP, GMP, ERP and BEM fates along increasing pseudotime (Fig. 2F). We observed smooth trends in expression and accessibility of important lineage markers along each fate trajectory, with *MPO*, *DNTT*, *GATA1* and *HDC* gene trends increasing along respective GMP, LMPP, ERP, and BEM fates (Fig. 2F,G, Fig. S2E). Interestingly, the trend in *GATA1* expression across erythroid and BEM development fits with recent descriptions of BEM as derived from the megakaryocyte-erythroid progenitors (*47–49*), characterized by a transient peak followed by attenuation of *GATA1* activity in BEM. Based on cell fate probabilities, we generated branch masks to isolate cells along lymphoid, myeloid, and erythroid lineages, and calculated frequency of cells from each infant timepoint along these branches (Fig. 2F,G). For all lineages, we observed a decreasing frequency of Day 4 cells over pseudotime (more HSC, fewer intermediate progenitors), and increasing frequencies of Month 3 and 13 cells (relatively fewer HSC and more intermediate progenitors) (Fig. 2G). Although the sample pooling and pre-enrichment of CD34^+^ cells prevents us from accurately quantifying cHSPC proportions out of total PBMC over development, we also noted an age-associated decrease in the abundance of circulating CD34^+^ progenitors (among Lineage^-^) during post-enrichment CD34^+^ cell sorting (Fig. S2F). Overall, our results support a shift in the cHSPC population over the first year of life from greater numbers of highly plastic, multipotent HSC (that includes unique B1 potential) to lower numbers of more differentiated progenitor subtypes.

## Consistent, age-associated trends in HSC gene expression

Given the observed shifts in cHSPC subtype frequency and heterogeneity within these subclusters across age, we next compared transcriptional phenotypes within the more homogenous population of undifferentiated HSCs, to better understand early life changes in stem cell states and differentiation bias. Differential gene expression results comparing infant to adult HSCs revealed many significant changes across all collected timepoints (Fig. 3A-C, Table S3). Day 4 upregulated genes included a number of markers of stemness and reconstitution potential, such as *HLF*, *INPP4B*, and *MLLT3* (Fig. 3A), supporting our observation that the majority of infant circulating progenitors at 4 days of age are enriched for more primitive, quiescent and/or self-renewing phenotypes (*50–52*). In contrast, upregulated genes at Month 3 revealed clear indications of NF-κB-associated inflammation and activation, with *NFKB1* as one of the most significant hits, followed by *REL*, *TANK*, *NFKBIA*, *CXCL8*, and *CD69* among others (Fig. 3B). Finally, Month 13 upregulated genes were characterized by interferon response and oxidative phosphorylation pathways, including interferon stimulated genes (*ISG15*, *IFI44L*, *IFITM2*), antigen presentation machinery (*HLA-A*, *HLA-B*, *HLA-C*), and electron transport chain components (*COX4I1*, *COX8A*, *NDUFA11*, *NDUFA4, NDUFC2*) (Fig. 3C). Given clear differences observed between each early-life timepoints and adulthood, we next asked whether there might be broader patterns in HSC gene expression during the first year of life that one-to-one comparisons might miss.

**Fig. 3.**
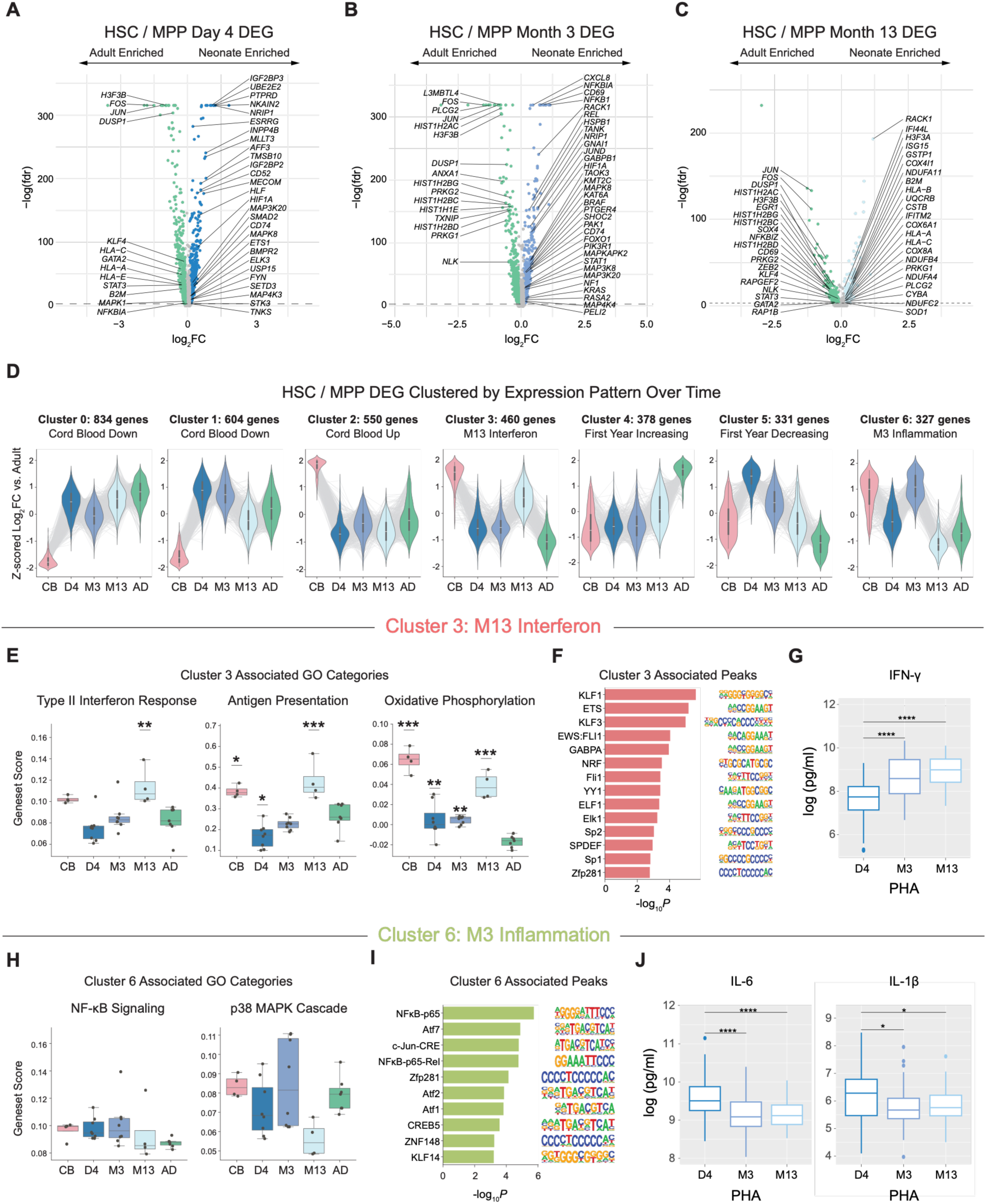
HSC gene expression follows consistent, age-associated programs. (**A-C**) Differential gene expression results in HSC/MPP comparing Day 4 (A), Month 3 (B) and Month 13 (C) to adult samples. Volcano plots show log_2_FC and -log(FDR) per gene, with genes with abs(log_2_FC) > 0.1 and FDR < 0.001 marked as significant. (**D**) Differential genes were clustered according to z-scored log_2_FC vs. adult to capture HSC / MPP gene trajectories over time. For each gene cluster, violin plots show z-scored log_2_FC distribution for all cluster genes per timepoint, with lines connecting individual genes across timepoints. (**E**) Box plots of per sample expression score in HSC/MPP of gene modules significantly enriched in Cluster 3 genes across timepoints. Expression scores were compared across timepoints by one-way ANOVA, with pairwise differences from the adult group determined by Tukey’s HSD test (adjusted P values shown). p > 0.05 = ns, ∗p ≤ 0.05, ∗∗p ≤ 0.01, ∗∗∗p ≤ 0.001. (**F**) Bar plot of HOMER motif enrichment results for snATAC peaks accessible in HSC / MPP at Cluster 3 gene loci, visualizing -log_10_*P* of enrichment and sequence Position Weight Matrix (PWM) per motif. (**G**) Log-normalized concentration (pg/mL) of IFN-γ upon 96h whole-blood stimulation with PHA of blood samples from n=66 control infant samples collected at 4 days, 3 months, and 13 months post-randomization. Statistical significance was assessed using a repeated-measures one-way ANOVA followed by paired post-hoc comparisons between timepoints with Holm correction for multiple testing. p > 0.05 = ns, ∗p ≤ 0.05, ∗∗p ≤ 0.01, ∗∗∗p ≤ 0.001. (**H**) Box plots of per sample expression score in HSC/MPP of gene modules significantly enriched in Cluster 6 genes across timepoints. Expression scores were compared across timepoints by one-way ANOVA, with pairwise differences from the adult group determined by Tukey’s HSD test (adjusted P values shown). (**I**) Bar plot of HOMER motif enrichment results for snATAC peaks accessible in HSC / MPP at Cluster 6 gene loci, visualizing -log_10_*P* of enrichment and sequence PWM per motif. (**J**) Log-normalized concentration (pg/mL) of IL-6 and IL-1β upon 24h whole-blood stimulation with PHA of blood samples from n=66 control infant samples collected at 4 days, 3 months, and 13 months post-randomization. Statistical significance was assessed using a repeated-measures one-way ANOVA followed by paired post-hoc comparisons between timepoints with Holm correction for multiple testing. p > 0.05 = ns, ∗p ≤ 0.05, ∗∗p ≤ 0.01, ∗∗∗p ≤ 0.001.

To describe transcriptional trends over time, we clustered genes according to their change in expression over all timepoints, resulting in a total of 7 HSC gene clusters (Fig. 3D, Table S4). When samples were scored by cluster gene expression, cluster-specific trends were highly consistent across individuals, indicating that hematopoiesis follows a structured pattern of transcriptional development over the first year of life (Fig. S3B). This analysis also highlighted the dramatic transcriptional differences between cord blood HSCs and all other timepoints, with Cluster 0, Cluster 1, and Cluster 2 largely driven by genes highly up- or down-regulated in cord blood. Given that cord blood has been used in the past as a rich source of HSCs in studies of hematopoiesis, we emphasize that cord blood HSCs in our data are not representative of the transcriptional landscape of infant HSCs, even only a few days after birth. While part of this difference may be explained by our cord blood and infant samples deriving from separate cohorts, previous studies have also observed significant changes in phenotype between cord blood and infant PBMC (*13, 53*). Differential gene expression analysis of cord blood HSCs and Day 4 HSCs in our data revealed large shifts in gene expression, with notable higher expression of cell cycle genes such as *CCNL1*, *CCND2*, and *CCNK* in cord blood, and lower expression of HSC quiescence-associated TFs *HLF* and *ERG* (Fig. S3A). Multiple groups have described fetal HSCs to have greater self-renewal and proliferative properties than adult HSCs (*45, 54, 55*). The cell-cycle signatures we see in cord blood in our data may indicate that cord blood, being collected at the time of birth, is a better representation of the late fetal stage than neonatal hematopoiesis.

Among the other gene clusters, we observed that Clusters 4 and 5 displayed a stepwise transition from infant to adult expression levels, with Cluster 4 genes progressively increasing from Day 4 to adults, and Cluster 5 genes progressively decreasing with age (Fig. 3D). Notable genes in Cluster 4 included lineage-specific transcription factors (*LMO2*, *NFATC1*, *GATA2*, *KLF4*, and *KLF9*), regulators of the balance between self-renewal and differentiation (*NFATC1, TET2*, *BCL2*), and DNA damage repair factors (*TP53BP1*, *RNF168, SMC4, SMC5, HMGB2*) (*56–58*). Activation of DNA repair pathways has been shown to be associated with HSC entry into the cell-cycle, to safeguard against accumulated DNA damage during long periods of quiescence (*59*). Thus, seeing DNA repair machinery in combination with lineage-specific factors in Cluster 4 is indicative of a program of increasing HSC lineage-priming and differentiation with age, which concurs with our HSPC subcluster abundance analysis (Fig. 2G). Cluster 5, on the other hand, contains core regulators of HSC maintenance and quiescence (*HLF*, *MLLT3*, *CD74*) and factors associated with lymphopoiesis, especially B-lymphopoiesis (*MEF2C*, *PTEN*, *IGHM*, *IKZF1*) (*50, 52, 60–63*). While stemness-related genes align with the higher abundance of true HSCs in infants relative to adults, the enrichment of B-lineage genes in Cluster 5 likely reflects the elevated B-cell proportion we observe in infant PBMC. Cluster 5 seems to capture the unique properties of post-natal hematopoiesis—balancing a high level of HSC plasticity with robust B-cell output. Overall, Clusters 4 and 5 define signatures of age-associated changes in HSC: capturing the transition from a unique, post-natal hematopoietic state enriched for regulators of the core HSC program and lymphopoiesis, to the mature adult phenotype of increased lineage specification and differentiation.

Finally, we found Cluster 3 and Cluster 6 to be largely infant-specific, with Cluster 3 expression elevated at the Month 13 timepoint, and Cluster 6 at Month 3 (Fig. 3D). Interestingly, Cluster 3 and Cluster 6 included genes associated with inflammation and immune response (*HLA-A*, *B2M*, and *ISG15* for Cluster 3, and *NFKB1*, *REL*, and *IRF1* for Cluster 6), which led us to explore these two gene clusters in more depth, although we provide gene cluster assignments and gene set enrichment results for all clusters as a resource (Table S4).

As HSC differentiate and exit quiescence, they shift from primarily glycolytic energy production to oxidative phosphorylation (*64*). Furthermore, both Type I and Type II interferons have been implicated in HSC loss of quiescence and entry into the cell-cycle in response to infections (*65, 66*). We found Cluster 3 genes (increased at Month 13) to be significantly enriched for the GO terms *aerobic electron transport chain (GO:0019646)*, *cellular response to type I interferon (GO:0071357)* and *cellular response to interferon-gamma (GO:0071346)*, among other terms related to oxidative phosphorylation and interferon response. Expression of genes in these GO terms followed the same pattern over timepoints as Cluster 3 genes as a whole (Fig. 3E). Accessible peaks in HSC surrounding cluster gene loci were enriched for KLF and ETS family motifs, many of which play important roles in hematopoietic cell fate, alongside NRF and YY1 family motifs, which are implicated in mitochondrial biogenesis and oxidative function (*67, 68*) (Fig. 3F). Overall, analysis of Cluster 3 genes and related TF motifs led us to conclude that this cluster captures increased HSC differentiation and an accompanying metabolic switch with increased oxidative phosphorylation at Month 13, alongside—and perhaps induced by— an interferon response.

At the same time, we noted that CD8^+^ Memory T Cells and NK cells, both of which are major interferon-producing cell types, peak in abundance in our dataset at 13 months of age (Fig. 1E). Therefore, we hypothesized that the activation of Cluster 3 gene expression at Month 13 in HSC may be a result of increased interferon production in the infant immune system as a whole, a feature likely driven by cumulative microbial exposure, associated memory T cell responses, and IFN-γ production. To validate functionally whether Cluster 3 gene expression corresponded with increasing IFN-γ production during the first year of life, we examined whole-blood cytokine production upon stimulation with phytohemagglutinin (PHA) or *Candida albicans* (*C. albicans*) in PBMC isolated from children in the Calmette cohort (*34*). Indeed, IFN-γ production capacity increased significantly from Day 4 to Month 13 timepoints (Fig. 3G, Fig. S3C), thereby linking IFN-γ increases with hematopoietic maturation in the first year of life.

We found Cluster 6 genes (increased at Month 3) to be significantly enriched for the GO terms *regulation of I-kappaB kinase/NF-kappaB signaling (GO:0043122)* and *regulation of p38MAPK cascade (GO:1900744),* with expression of NF-κB pathway genes being upregulated in early infant timepoints as compared to adults (Fig. 3H). Motif enrichment of associated peaks revealed a clear NFκB-p65 signal, alongside AP-1 and ATF activity (Fig. 3I). Given the link between Cluster 6 genes and NF-κB signaling, we hypothesized that this cluster may represent an overall HSC response to inflammation, perhaps triggered upon exposure to microbial products and stress associated with birth in the first days of life. We therefore examined IL-6 and IL-1β production among PBMC in response to PHA or *C. albicans* (*34*), given that both of these key inflammatory cytokines are induced through NF-κB pathway activation. We found that IL-6 and IL-1β production in PBMC decreased from Day 4 to Month 3 and Month 13 timepoints, consistent with an overall downward trend in inflammatory signaling encoded at the level of HSC (Fig. 3J, Fig. S3D), and linking NF-κB activity in HSPC with hyper-inflammatory responses in early life.

Overall, analysis of Cluster 3 and Cluster 6 gene expression revealed common trends in infant hematopoietic and immune development in the first year of life. HSC transcriptomic analysis revealed a shift from early inflammation programs prominently featuring NF-κB pathway activity through three months—supported by increased production of IL-6 and IL-1β in mature immune cells—to a maturation program featuring higher expression of IFN-γ-response genes by 13 months.

## Early activation of interferon responses in infant HSPC associated with BCG vaccination

Having established gene expression programs in control infant HSC in our data, we subsequently assessed the impact of BCG vaccination on those programs. While we did not observe significant effects of BCG vaccination on HSPC subtype abundance (Fig. S4A), PCA analysis of pseudo-bulk RNA data from infant HSC revealed clear and consistent impacts of BCG on gene expression, which varied according to the sample timepoint (Fig. 4A). We then asked whether these transcriptional effects represented changes in HSC gene clusters established in the previous section and found significant changes in the majority of gene clusters, but especially in clusters 1 and 3 (Fig. 4B, Fig. S4B). For cluster 3, which we previously described as reflecting an increased IFN-γ signature at 13 months of age, we saw an increase in cluster gene expression in BCG samples relative to control at Month 3. We compared BCG Month 3 vs. control Month 3 upregulated genes (Fig. 4C) to the control group HSC Cluster 3 gene set that increases through the first year, given its association with interferon response, and found significant intersection (Fig. 4D). Consistent with this and Cluster 3 enrichment reflecting an IFN-γ response, BCG upregulated genes at Month 3 included antigen presentation machinery (*HLA-A*, *HLA-C*, *HLA-E*, *B2M*) and electron transport chain components (*NDUFA4*, *NDUFS5*, *COX4I1*, *UQCRB*) (Fig. 4C,G, Table S5). This is consistent with previously described effects of BCG vaccination on the hematopoietic compartment in mouse and adult human studies, which report an increase in IFN-γ and oxidative phosphorylation programs in HSC for 1 to 3 months following vaccination (*25, 27*). To compare the effects of BCG on HSPC in adults and children, we compared BCG upregulated genes at Month 3 in HSPC from our neonatal dataset to the set of BCG immunotherapy upregulated genes in adult cHSPC from a cohort of bladder cancer patients reported by Daman et al. 2025 (*28*), and again found significant enrichment (Fig. 4E). These results indicate that the transcriptional programs in progenitor cells upregulated 3 months post-BCG vaccination in neonates resemble not only the programs associated with BCG treatment in adults, but also a common IFN-γ signature that in control samples appears later in HSC development—at 13 months.

**Fig. 4.**
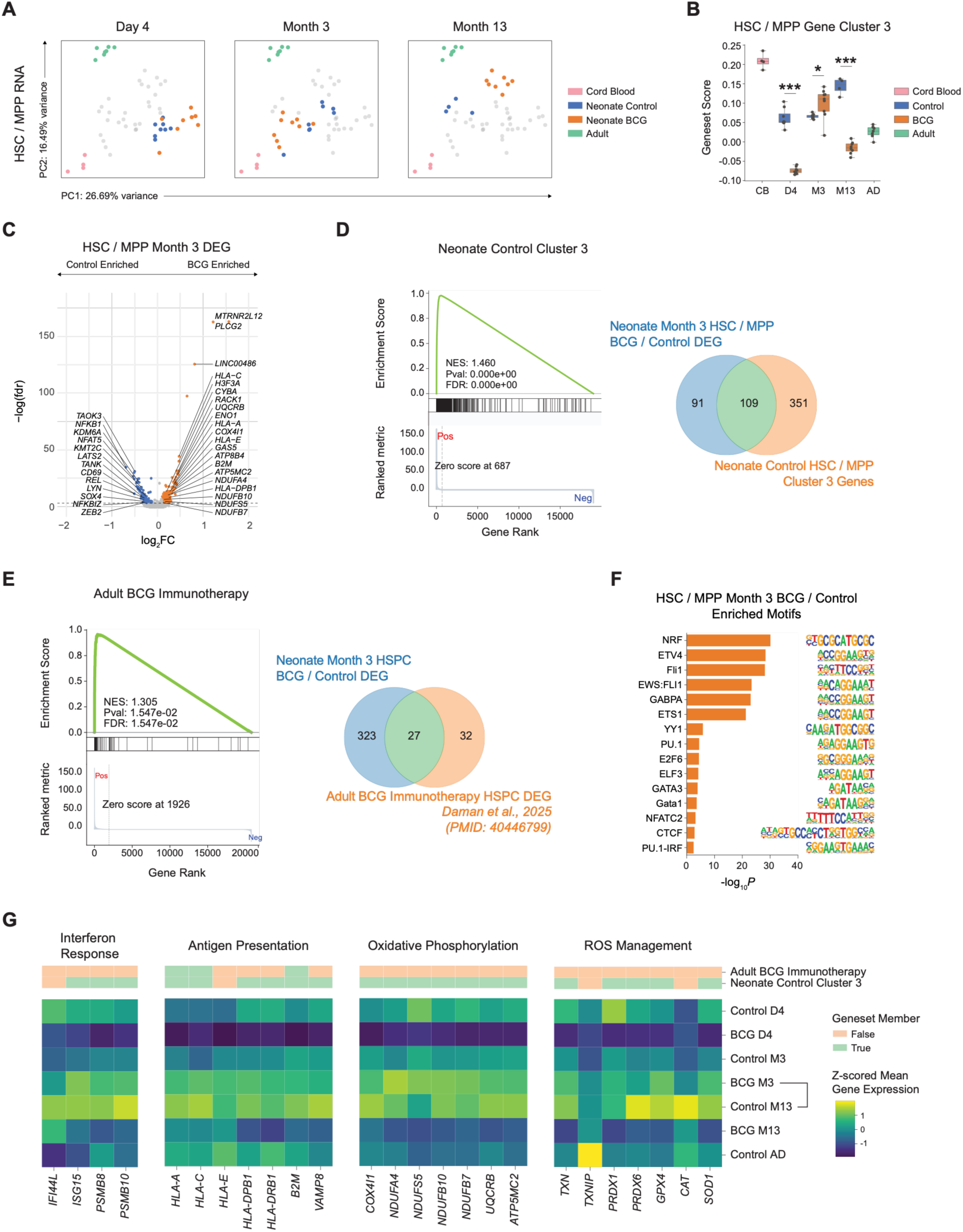
BCG vaccination accelerates expression of interferon response programs in HSC. (**A**) PCA plot of pseudo-bulk HSC /MPP snRNA data, highlighting samples from Day 4, Month 3, and Month 13 infant timepoints. (**B**) Box plots of per sample expression score in HSC/MPP of Cluster 3 genes as defined in Fig. 3. Differences between BCG and control groups at each timepoint were assessed using an unpaired two-sided Student’s *t*-test, with *p*-values adjusted for multiple comparisons using the Holm method. p > 0.05 = ns, ∗p ≤ 0.05, ∗∗p ≤ 0.01, ∗∗∗p ≤ 0.001. (**C**) Differential gene expression results in HSC/MPP comparing BCG to control samples at Month 3. Volcano plot shows log_2_FC and -log(FDR) per gene, with genes with abs(log_2_FC) > 0.1 and FDR < 0.001 marked as significant. (**D**) Gene set enrichment analysis (GSEA) results comparing Month 3 BCG vs. control differential gene expression in HSC / MPP to the Cluster 3 geneset (left), and a Venn diagram visualizing the number of shared genes (right). (**E**) GSEA results comparing Month 3 BCG vs. control differential gene expression over all HSPC to the Daman et al. 2025 adult BCG immunotherapy gene signature (left), and a Venn diagram visualizing the number of shared genes (right). (**F**) Bar plot of HOMER motif enrichment results for upregulated peaks within Month 3 BCG vs. control differential peaks in pseudo-bulk HSPC snATAC data. Plot visualizes -log_10_P of enrichment and sequence PWM per motif. (**G**) Heatmap of mean gene expression for selected significant genes (columns) from the DEG in (C), z-scored across conditions and grouped according to biological function. Heatmap annotations indicate gene membership in the adult BCG immunotherapy or control cluster 3 genesets shown in (D) and (E).

We then looked at BCG-associated changes in HSC at Month 3 in more detail. In differential peak analysis of snATAC-Seq chromatin accessibility data comparing BCG and control HSC at Month 3 (Table S6), we found upregulated peaks to be significantly enriched for multiple ETS family motifs, alongside NRF and YY1 (Fig. 4F). These motifs matched the HSC Cluster 3 gene set-associated motifs previously described, which peak at Month 13 in BCG-naive infants over the first year of life, and are often associated with expansion and associated metabolic changes. To validate the link between BCG vaccination and interferon signaling in our PIE-Seq dataset, we examined IFN-γ protein concentrations in serum collected from the larger Calmette cohort. Serum IFN-γ concentrations showed a similar overall trajectory across infancy, rising from Day 4 to Month 13 in controls, and were directionally consistent with the HSC interferon-response signature; however, differences between BCG and control groups at individual timepoints were not statistically significant in this dataset (Fig. S4C).

Overall, examination of gene cluster expression indicated HSC induction of an earlier interferon response following BCG vaccination: BCG induced an HSC program characterized by oxidative phosphorylation and interferon-stimulated genes at 3 months that was not observed in control infants until 13 months of age. We found that BCG vaccination induced programs of interferon response, antigen presentation, and oxidative phosphorylation in HSC at the Month 3 timepoint (Fig. S4D), which echoed the adult response to BCG (*27, 28*). These programs are also a part of normal hematopoietic development in infants, where they become progressively more active in HSC towards 13 months of age. To summarize these signatures, we visualized expression of key genes within the set of significant BCG-upregulated genes at Month 3 in HSC over time. Across genes characteristic of interferon response, antigen presentation, and oxidative phosphorylation—in addition to reactive oxygen species (ROS) management, by which cells safely process the byproducts of oxidative phosphorylation—we saw a repeated pattern in which expression levels in the BCG Month 3 condition closely resembled Month 13 controls (Fig. 4G).

## An NRF1-associated program in HSC at Day 4 post-BCG vaccination

Previous studies of BCG vaccination in humans have not studied the early effects of BCG on hematopoiesis, but examination of early post-vaccination timepoints could provide key information on the molecular pathways that induce BCG’s long-term effects: this motivated us to assess the Day 4 timepoint in more depth. Differential gene expression results (Table S5) revealed hallmarks of cell division and exit from quiescence, with BCG-upregulated genes at Day 4 in HSC including cell cycle regulators (*CCND3*, *CCNL1*, *CDK12*) and genes associated with DNA repair (*ATM*, *TOPBP1*, *PRKDC*) (Fig. 5A). Significantly enriched GO terms included *G2/M transition of mitotic cell cycle (GO:0000086)* and *DNA repair (GO:0006281)* (Fig. 5B). We also saw enrichment of terms such as *protein ubiquitination (GO:0016567)* related to the ubiquitin-proteasome system, which has been linked to the regulation of HSC quiescence and activation (*69, 70*). Supporting induction of a program of HSC expansion and activation, we observed an increased proportion of CD34^+^ progenitors in peripheral blood during cell sorting for PIE-Seq at Day 4 (Fig. S4E). In contrast, genes dampened by BCG at Day 4 included those involved in antigen presentation (*HLA-A*, *HLA-B*, *HLA-C*, *B2M*) and oxidative phosphorylation (*UQCRB*, *COX7C*, *COX4I1*) (Fig. 5A,B), corresponding to the HSC Cluster 3 gene program— the only cluster downregulated by BCG at this early timepoint (Fig. S4B,D).

**Fig. 5.**
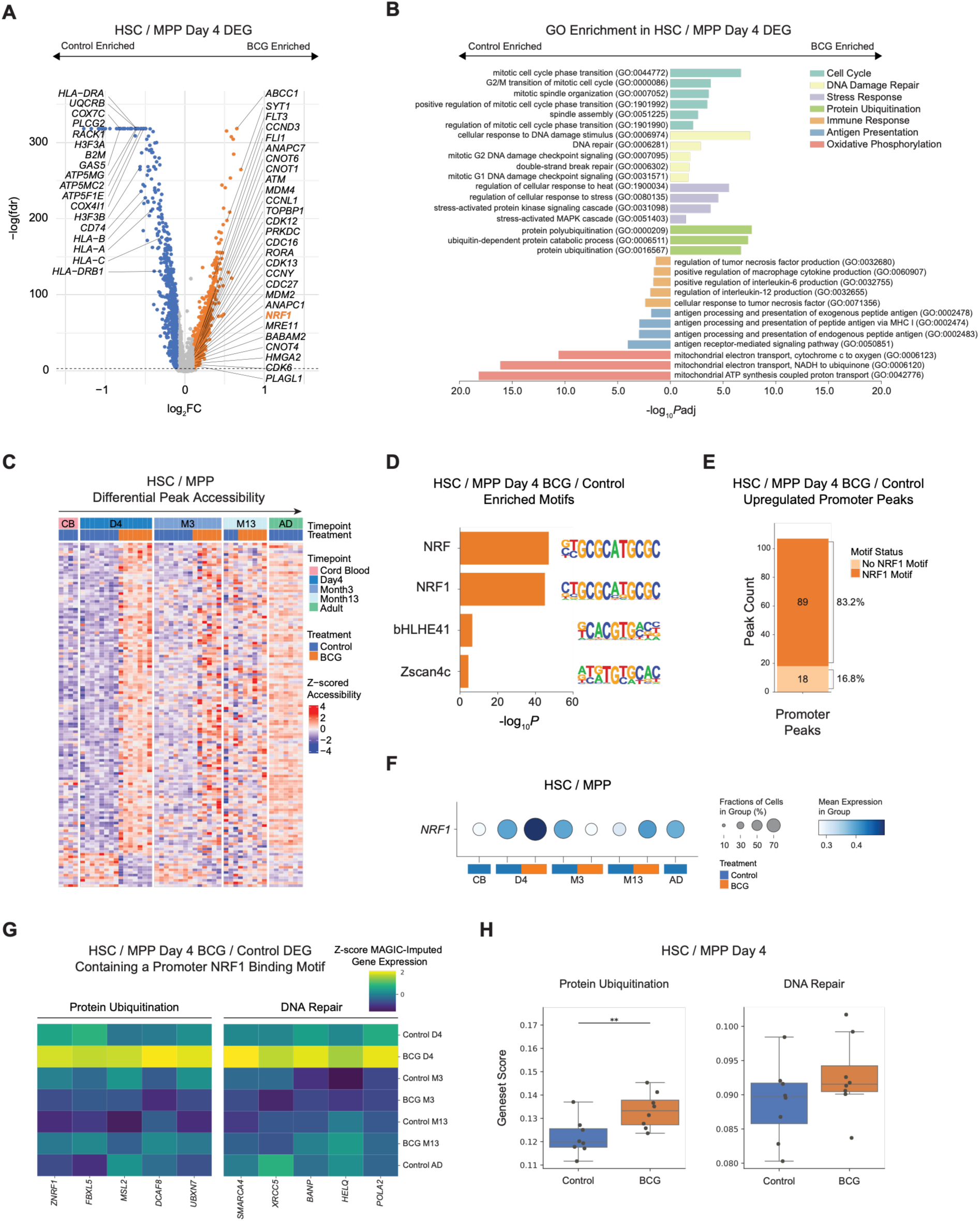
NRF1 drives early HSC response to BCG-vaccination. (**A**) Differential gene expression results in HSC/MPP comparing BCG to control samples at Day 4. Volcano plot shows log_2_FC and -log(FDR) per gene, with genes with abs(log_2_FC) > 0.1 and FDR < 0.001 marked as significant. (**B**) Bar plot visualizing -log_10_Padj of GO enrichment for selected terms enriched in significant Day 4 BCG vs. control upregulated (right) and downregulated (left) genes. (**C**) Heatmap of z-scored accessibility at Day 4 BCG vs. control differential peaks (rows) in pseudo-bulk HSPC snATAC data across timepoints and conditions (columns). (**D**) Bar plot of HOMER motif enrichment results for upregulated Day 4 peaks in (C). Plot visualizes -log_10_P of enrichment and sequence PWM per motif. (**E**) Bar plot of proportion of promoter-annotated peaks containing an NRF1 motif peak in the set of peaks significantly more accessible in the BCG vs. Control Day 4 comparison. (**F**) Dot-plot of *NRF1* expression in HSC/MPP. Dot size represents the number of cells per group expressing NRF1, and dot color represents log-normalized expression level. (**G**) Candidate NRF1-regulated genes were selected as genes with promoter peaks containing an NRF1 motif, with a significant increase in promoter accessibility in the BCG vs. Control Day 4 comparison. Heatmap shows z-scored mean expression of selected genes, using MAGIC-imputed expression values. (**H**) Box plots of per sample expression score in Day 4 HSC/MPP of GO terms selected based on candidate NRF1-regulated genes. Differences between BCG and control groups at each timepoint were assessed using an unpaired two-sided Student’s *t*-test. p > 0.05 = ns, ∗p ≤ 0.05, ∗∗p ≤ 0.01, ∗∗∗p ≤ 0.001.

To uncover possible regulators of HSC activation at Day 4, we looked at differential peaks between BCG and control samples in HSC (Table S6). Visualization of the accessibility of these peaks across all collected timepoints revealed an interesting pattern suggestive of memory characteristics: BCG-upregulated peaks at Day 4 remained open relative to control samples, and downregulated peaks remained closed (Fig. 5C). Furthermore, the BCG-induced chromatin state at these peaks at Day 4 resembled adult controls far more than other infant samples. To identify transcription factors driving these changes, we examined enriched motifs in the regions gaining in accessibility. We found upregulated peaks at Day 4 to be dominated by NRF/NRF1 activity, with more than 83.2% of differential promoter regions gaining in accessibility containing an NRF motif (Fig. 5D,E). We also saw upregulation of *NRF1* in our Day 4 differential gene expression results (Fig. 5A,F), which increased our confidence in identifying NRF1 as a major factor of interest in early BCG response. NRF1 is a master regulator of mitochondrial biogenesis, with a binding site at the promoters of multiple respiratory complex subunits, as well as factors involved in mtDNA transcription and replication (*71*). In murine HSC, elevated NRF1 activity upon knockdown of its repressor SIRT7 has been linked to increased mitochondrial counts and proliferation under homeostatic conditions (*72*). In addition, bulk ATAC-sequencing of adult human PBMC post-BCG vaccination identified enriched NRF1 motif accessibility among strong trained-immunity responders (*73*). Consistent with our observation of persisting chromatin accessibility at NFR1 motif-enriched sites following BCG, NRF1 was also recently described among a select set of methyl-sensitive TFs whose activity is associated with long-term chromatin memory (*74*).Together, our findings highlight NRF1 as a compelling candidate transcription factor mediating early chromatin remodeling, coupled to metabolic adaptation during the early HSC response to BCG.

In order to evaluate the downstream effects of NRF1 activity in HSC, we compiled a list of candidate NRF1 regulated genes according to two criteria: first, that the gene promoter contained an NRF1 motif, and second, that the gene promoter peak gained in accessibility in the BCG vs. Control Day 4 comparison in HSC. This constituted a total of 74 genes expressed in our snRNA data, 52 of which were significantly up or down-regulated (p < 0.05 from MAST results in Day 4 BCG vs. control HSC) in expression in HSC at Day 4 (Fig. S4F). While the majority of these targets were significantly upregulated, a number were down-regulated, indicating that NRF1 binding may act to both activate or repress transcription depending on context. Notable BCG Day 4-upregulated genes among these putative NRF1 targets included components of the ubiquitin-proteasome system (*ZNRF1, FBXL5, MSL2, DCAF8, UBXN7*) and DNA damage repair factors (*SMARCA4, XRCC5, BANP, HELQ, POLA2*) (Fig. 5G). These target pathways correspond to those previously reported in NRF1 ChIP-Seq data in human neuroblastoma cells (*75*). Furthermore, scoring of HSC according to GO terms *protein ubiquitination (GO:0016567)* and *DNA repair (GO:0006281)* demonstrated consistent upregulation of these pathways in HSC for BCG samples at the Day 4 timepoint (Fig. 5H).

In conclusion, BCG vaccination is associated with a rapid HSC expansion phenotype 4 days post-vaccination. This is accompanied by increased accessibility in a number of genomic regions, the majority of which contain NRF1 motifs, that then remain open for all observed timepoints even while the activated HSC gene expression program resolves. Analysis of candidate NRF1 target genes at this timepoint suggest that NRF1 activation marks a coordinated program in HSC in which cells move towards cell cycle entry, while simultaneously upregulating DNA damage repair and protein ubiquitination pathways to handle the stress and increased protein turnover associated with exit from quiescence.

## A population of infant myeloid cells in a highly inflammatory alternative state featuring increased NFκB activity (MyNFKB1^hi^)

Our analyses of circulating HSPC demonstrate structural shifts in composition and transcriptional profile over the first year of life. To explore how shifts in the HSPC population might impact differentiated progeny cells, we next examined mature myeloid cells in both infants and adults. Upon visualization of snATAC embeddings of the myeloid compartment, we were surprised to see that each major myeloid cell type, as defined by marker gene expression of *CD14* (CD14^+^ Monocytes), *FCGR3A* (CD16^+^ Monocytes), and *HLA-DRA* (cDC), could be separated into two mirrored groups of cell-type clusters which were largely defined by either early neonatal or Month 13/Adult origin (Fig. 6A-D, Fig. S5A). Using diffusion maps to characterize major axes of epigenetic variation (*76, 77*), we found that the primary axis (Diffusion Component 1) clearly separated each cell type into two distinct subsets, matching the two mirrored cluster groups observed in the embedding (Fig. 6A). In addition, Diffusion Component 1 was strongly correlated with key markers of inflammation and activation including *NFKB1*, *IL1B*, and *NLRP3* (Fig. 6E, Fig. S5E). Therefore, to better characterize this programmatic axis of myeloid variation, we annotated myeloid cell clusters low in Diffusion Component 1 as Myeloid Conventional (MyConv) and clusters high in Diffusion Component 1 as Myeloid NFKB-high (MyNFKB1^hi^) (Fig. 6C).

**Fig. 6.**
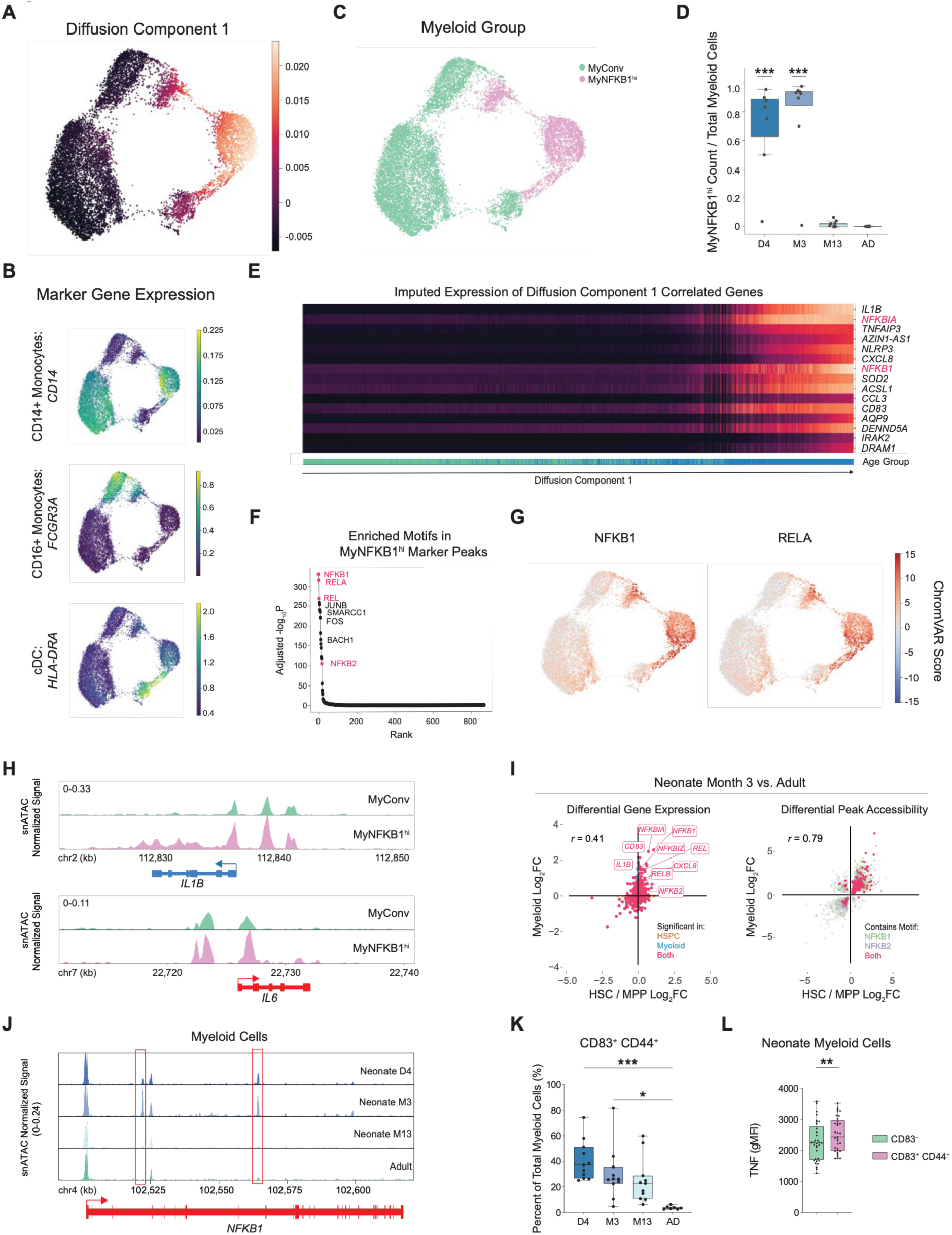
An NFκB-driven inflammatory myeloid cell state (MyNFKB1^hi^) in infants. (**A**) UMAP visualization of snATAC data from myeloid cells in all samples, colored by Diffusion Component 1 value. (**B**) UMAP of myeloid cells as in (A) colored by MAGIC-imputed expression of myeloid subtype marker genes. (**C**) UMAP of myeloid cells as in (A) colored by myeloid group annotation. (**D**) Box plot of proportion of MyNFKB1^hi^ cells out of total myeloid cells per sample across all control timepoints. Proportions were compared across timepoints by one-way ANOVA, with pairwise differences from the adult group determined by Tukey’s HSD test (adjusted P values shown). p > 0.05 = ns, ∗p ≤ 0.05, ∗∗p ≤ 0.01, ∗∗∗p ≤ 0.001. (**E**) MAGIC-imputed expression of Diffusion Component 1 correlated genes (rows) across all myeloid cells (columns). Cells are ordered by Diffusion Component 1, and annotated by age group at bottom. (**F**) Rank plot of motif enrichment results in marker peaks for MyNFKB1^hi^ vs. MyConv cells, with -log_10_P-adjusted on the y-axis and rank on the x-axis for each motif. (**G**) UMAP of myeloid cells as in (A) colored by chromVAR score per cell for NFKB1 and RELA motifs. (**H**) Pseudo-bulk ATAC-seq accessibility tracks of the *IL1B* and *IL6* loci for MyConv and MyNFKB1^hi^ cells. (**I**) Scatter plots of gene expression Log_2_FC (left) and pseudo-bulk ATAC peak accessibility Log_2_FC (right) for control Month 3 vs. adult HSC / MPP (x-axis) and myeloid cells (y-axis). Genes are colored by cell-type significance, and peaks are colored by whether they contain an NFκB-family motif. Pearson correlation coefficient for genes or peaks significant in both HSPC and myeloid cells reported on plot. (**J**) Pseudo-bulk ATAC-seq accessibility tracks of the *NFKB1* locus across control D4, M3, M13, and adult timepoints. (**K**) Box plot depicting percentage of CD83^+^CD44^+^ cells out of total myeloid cells across control timepoints and adult samples, determined by flow cytometric analysis. Percentages were compared across timepoints by one-way ANOVA, with pairwise differences from the adult group determined by Tukey’s HSD test (adjusted P values shown). p > 0.05 = ns, ∗p ≤ 0.05, ∗∗p ≤ 0.01, ∗∗∗p ≤ 0.001. (**L**) Box plot of geometric mean fluorescence intensity (gMFI) of TNF in CD83^+^CD44^+^ myeloid cells and non-CD83^+^CD44^+^ myeloid cells in neonate samples. *p* values were calculated using a Wilcoxon matched-pairs rank sum test. p > 0.05 = ns, ∗p ≤ 0.05, ∗∗p ≤ 0.01, ∗∗∗p ≤ 0.001.

These two states were not only observed in snATAC visualizations, but also in snRNA, indicating strong transcriptional as well as epigenetic differences between them (Fig. S5B). Quantifying based on these annotations, we saw that MyNFKB1^hi^ cells were highly associated with infant samples, with the majority of myeloid cells from control infant Day 4 and Month 3 timepoints being classified as MyNFKB1^hi^ (Fig. 6D). Given the strong age-group association of MyNFKB1^hi^ cell state, we conducted multiple analyses to rule out the possibility of batch-effects driving observed myeloid variance. Visualization of MyNFKB1^hi^ vs. MyConv proportion per sequencing lane confirmed that these annotations were not associated with specific sample pools: the majority of batches contained both myeloid groups (Fig. S5C). Returning to the MrVI model trained on the full dataset, we also evaluated MrVI sample distances over all myeloid cells: this distance matrix measures how the sample of origin affects a cell’s representation in the sample-aware *z* latent space. Importantly, inclusion of batch as ‘nuisance’ covariate during model training should ensure that the *z* latent is minimally affected by batch and instead captures biological differences between samples. Hierarchical clustering of MrVI sample distances grouped Day 4 and Month 3 samples separately from other timepoints, supporting that the distinct myeloid phenotype observed in these early timepoints is driven by biological differences between samples (Fig. S5D).

To investigate biological drivers of this variation, we conducted differential gene expression analysis between MyConv and MyNFKB1^hi^ clusters, confirming upregulated expression of the inflammatory markers identified by correlation analysis in the MyNFKB1^hi^ subset (Table S7). Marker genes include NF-κB subunits *NFKB1* and *REL*, pathway regulators *NFKBIA*, *NFKBIZ*, *TNFAIP3*, and *TNIP1*, the *NLRP3* inflammasome sensor and its downstream effector *IL1B*, and NF-κB target cytokines and chemokines including *TNF*, *CXCL8*, *CCL3*, and *CCL4.* These markers are consistent with coordinated induction of the myeloid IL-1β pathway, where NF-κB-mediated signals induce transcription of pro-IL-1β, while the NLRP3 inflammasome processes it into bioactive IL-1β and primes myeloid cells for rapid inflammatory cytokine release (*78*). Analysis of MyNFKB1^hi^ marker peaks underscored the importance of NF-κB, with the top three enriched motifs all belonging to the NF-κB family (Fig. 6 F,G, Table S7). Finally, pseudo-bulk ATAC-Seq tracks from MyConv and MyNFKB1^hi^ cells revealed increased accessibility at the promoters and along the gene body of the key myeloid inflammatory cytokine genes *IL1B* and *IL6* (Fig. 6H). Overall, the MyNFKB1^hi^ phenotype was defined by high NF-κB activity, as evident in transcription and accessibility of both *NFKB1* and its downstream targets.

Given our findings of NF-κB activity in early infant timepoints in HSC (Fig. 3H,I), we questioned whether inflammatory programs in HSC and mature myeloid cells might reflect a shared regulatory circuit transmitted from progenitors to their progeny. To compare these compartments, we plotted the fold change of significantly differentially expressed genes in control Month 3 infants versus adults for HSCs, against the corresponding fold change in myeloid cells. Despite comparing two distinct lineages, gene expression programs showed moderate correlation (Pearson’s R = 0.41), and *NFKB1* stood out as a top upregulated gene in both populations (Fig. 6I). Performing the same analysis on differential chromatin accessibility revealed a much stronger correlation (Pearson’s R = 0.79), particularly within peaks containing NFKB1 or NFKB2 motifs (Fig. 6I). This strong correlation in accessibility suggests that NF-κB target loci are broadly primed at the chromatin level in both progenitors and descendants. The prominence of NF-κB activity in reshaping chromatin state led us to examine the regulation of the *NFKB1* locus itself in infant samples. Visualization of *NFKB1* locus accessibility in myeloid cells revealed multiple infant-specific peaks that are absent by 13 Months and in adults (Fig. 6J). These findings support a model in which early-life regulation of *NFKB1* creates a developmental window of prominent NF-κB activity in chromatin across hematopoietic progenitors and the myeloid lineage, which results in a downstream inflammatory program that potentially influences myeloid cell responsiveness. Recent demonstration of NF-κB’s potent activity in *de novo* chromatin opening at inflammatory genes further highlights the potential for this pathway to durably reprogram HSPC and progeny myeloid cells (*79*).

We adopted both *in-silico* and experimental approaches to validate our observation of the MyNFKB1^hi^ cell state. To confirm that MyNFKB1^hi^ transcriptional programs exist in other infant datasets, we turned to scRNA sequencing data collected by Nouri et al. examining the response to routine vaccinations in PBMC of two-month old infants (*80*). We scored pre-vaccination myeloid cells from this dataset and control adult PBMC data according to a module of MyNFKB1^hi^ marker genes, and indeed identified higher expression of MyNFKB1^hi^ markers in infant samples (Fig. S5F). This finding confirms that the MyNFKB1^hi^ transcriptional program is identifiable in early-life myeloid cells, in a separate cohort with gene expression data collected using a different method.

We next aimed to validate the presence of MyNFKB1^hi^ cells in infants through flow cytometry. For experimental validation, we identified CD44 and CD83 as candidate surface markers for MyNFKB1^hi^ cells given that their respective coding genes were highly ranked markers of MyNFKB1^hi^ cell state in differential gene expression analysis of the Multiome data (Table S7), with *CD44* and *CD83* co-expression markedly enriched in MyNFKB1^hi^ compared to MyConv cells (Fig. S6A). We then developed a flow cytometry panel targeting these markers alongside canonical myeloid surface proteins with which to assess additional, previously untested Calmette cohort PBMC samples. While we found CD44 to be uniformly highly expressed on the surface of myeloid cells (CD45^+^ CD11b^+^ HLA-DR^+^), we saw myeloid cells with high surface CD83 staining almost exclusively in neonatal samples (Fig. S6B). We quantified the proportion of CD83^+^CD44^+^ cells out of total myeloid cells over all control timepoints and saw a clear decrease from Day 4 to Month 13 samples, with CD83^+^CD44^+^ cells being very rare in adults (Fig. 6K).

We then assessed the functionality of CD83^+^CD44^+^ cells through intracellular cytokine staining of TNF, IL-1β and IL-6. We first confirmed that myeloid cells in our samples were the dominant source of these cytokines, which supports the role of these proteins as key signaling molecules in myeloid-driven inflammation (Fig. S6C). We then compared cytokine concentrations in the CD83^+^CD44^+^ population to CD83^-^myeloid cells and saw significantly higher levels of intracellular TNF in CD83^+^CD44^+^ cells at unstimulated baseline conditions (Fig. 6L). Given the scarcity of infant samples, and the existence of a small proportion of CD83^+^CD44^+^ cells in adult samples, we used adult PBMC to profile the response of these cells to stimulation. Across stimulation with established activators of NF-κB signaling, including respiratory syncytial virus strain A2 (RSV-A2), *S. pneumoniae*, Resiquimod (R848), and Lipopolysaccharides (LPS), CD83⁺CD44⁺ myeloid cells exhibited robust, stimulus-dependent increases in intracellular TNF, IL-1β, and IL-6, whereas cytokine induction in CD83⁻ cells remained modest across conditions (Fig. S6D). These findings are consistent with enhanced NF-κB–associated inflammatory responsiveness in the CD83⁺CD44⁺ population. Given the concordance between marker expression, trends in abundance, and inflammatory cytokine expression, we suggest that the CD83^+^CD44^+^ monocyte population approximates the MyNFKB1^hi^ cell state observed in transcriptional and epigenetic data. Overall, these findings support a model of a transient early neonatal developmental window with a prominent (majority) myeloid program featuring elevated NF-κB activity, CD83 surface expression, enhanced signaling activity, and increased target gene expression, including heightened baseline and hyper-induced inflammatory cytokine production in response to bacterial or viral ligands.

## Transient downregulation of MyNFKB1^hi^ programs with BCG vaccination

Having established the MyNFKB1^hi^ phenotype in control samples, we next examined the effects of BCG vaccination on the myeloid compartment. While BCG vaccination has long been associated with enhanced responsiveness against heterologous immune challenges, it has also been reported that BCG downregulates low-grade systemic inflammation (*81, 82*). Any relation of such activities to progenitor or specific molecular myeloid programs is unknown. In our data, we identified a significant BCG-associated reduction in MyNFKB1^hi^ proportion out of total myeloid cells at 4 days post-vaccination, as well as a downward trend at 3 months (Fig. 7A). As an orthogonal approach, we assessed CD83^+^CD44^+^ myeloid cell abundance at Day 4 in BCG-vaccinated and control infants using additional Calmette cohort samples with the flow cytometry panel described above. Validating our findings in the multi-omics data, we saw that BCG indeed reduces the abundance of CD83^+^CD44^+^ myeloid cells at 4 days post vaccination (Fig. 7B).

**Fig. 7.**
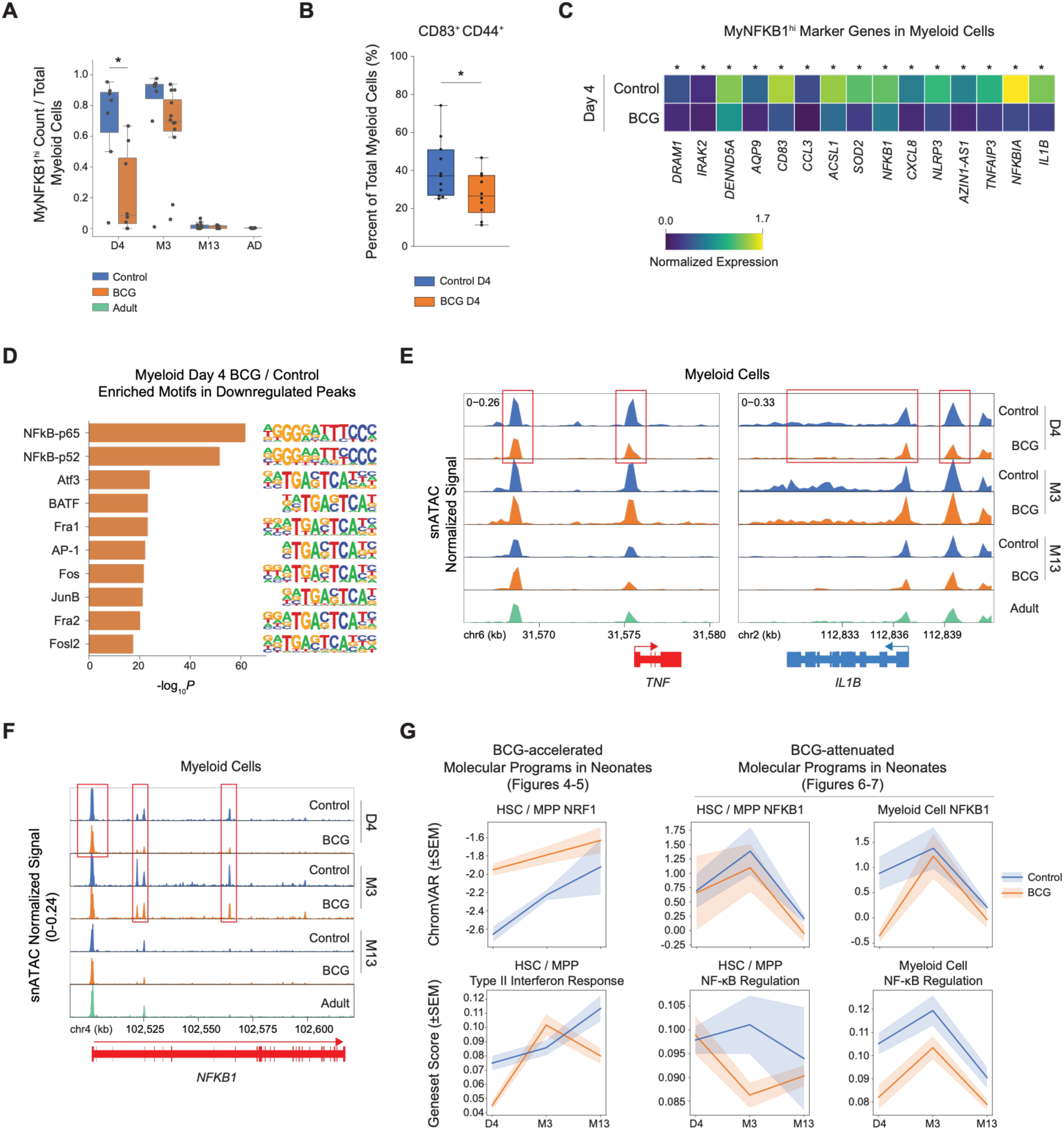
BCG vaccination temporarily attenuates infant myeloid inflammation. (**A**) Box plot of proportion of MyNFKB1^hi^ cells out of total myeloid cells per sample across all conditions and timepoints. Differences between BCG and control groups at each timepoint were assessed using an unpaired two-sided Student’s *t*-test, with *p*-values adjusted for multiple comparisons using the Holm method. p > 0.05 = ns, ∗p ≤ 0.05, ∗∗p ≤ 0.01, ∗∗∗p ≤ 0.001. (**B**) Box plot depicting percentage of CD83^+^CD44^+^ cells out of total myeloid cells at Day 4 for control and BCG neonate samples, as determined by flow cytometric analysis. *p* values were derived from an unpaired two-sided Student’s *t*-test. p > 0.05 = ns, ∗p ≤ 0.05, ∗∗p ≤ 0.01, ∗∗∗p ≤ 0.001. (**C**) Heatmap of mean log-normalized gene expression of MyNFKB1^hi^ marker genes (columns) in myeloid cells by condition (rows) at Day 4. A ∗ annotation indicates genes with p ≤ 0.001 and abs(log_2_FC) > 0.1 from MAST differential expression results. (**D**) Bar plot of HOMER motif enrichment results for downregulated peaks within Day 4 BCG vs. control differential peaks in pseudo-bulk myeloid snATAC data. Plot visualizes -log_10_P of enrichment and sequence PWM per motif. (**E**) Pseudo-bulk ATAC-seq accessibility tracks of the *TNF* and *IL1B* loci across timepoints and conditions in myeloid cells. (**F**) Pseudo-bulk ATAC-seq accessibility tracks of the *NFKB1* locus across timepoints and conditions in myeloid cells. (**G**) Summary of key observed phenotypes over time. Top row visualizes mean ChromVAR accessibility score of NRF1 and NFKB1 binding motifs for BCG and control groups in HSC/MPP and myeloid cells, ± standard error of the mean (SEM). The bottom row visualizes the mean expression score of key genesets ±SEM.

We subsequently assessed the functional consequences of BCG-induced decreases in MyNFKB1^hi^ abundance by evaluating the relationship between MyNFKB1^hi^ abundance and cytokine production in peripheral immune cells, by comparing MyNFKB1^hi^ proportion per individual to whole-blood stimulation data. At the month 3 timepoint, we saw a consistent positive correlation in control sample PBMCs from BCG-unvaccinated children between MyNFKB1^hi^ proportion and the production of key myeloid inflammatory cytokines IL-1β, TNF, and IL-6; conversely, we saw a consistent negative correlation with the production of the lymphoid-derived cytokines IFN-γ, IL-10, and IL-22 (Fig. S7A). We did not see a similar pattern in analysis of other timepoints. In BCG-vaccinated individuals, this correlation was lost, suggesting that BCG attenuates the relationship between MyNFKB1^hi^ cell state and inflammatory cytokine production over time (Fig. S7A).

To evaluate the molecular mechanisms underlying BCG-vaccination associated changes in myeloid phenotype, we examined transcriptional and epigenetic differences between BCG-experienced and control myeloid cells at Day 4 (Table S8). Changes in MyNFKB1^hi^ abundance were reflected in differential expression of key inflammatory genes across the total myeloid compartment, with significant downregulated genes at Day 4 including defining MyNFKB1^hi^ markers *NFKB1*, *NLRP3*, *IL1B*, and *CD83* (Fig. S7B). Visualizing expression of the top 15 MyNFKB1^hi^ markers in all myeloid cells at Day 4 showed consistent downregulation of expression post-BCG vaccination, especially in CD14^+^ monocytes, and with *CD83*, *NLRP3*, and *IL1B* among a set of consistently downregulated genes in the BCG condition across myeloid cell subtypes (Fig. 7C, Fig. S7C). Differential peak analysis across all myeloid cells at Day 4 revealed a total of 1,017 significant peaks, with the top motifs enriched in downregulated peaks being NFκB-p65 and NFκB-p52 (Fig. 7D, Fig. S7D). In examining pseudo-bulk ATAC accessibility at the gene loci of key myeloid cytokines *TNF* and *IL1B*, we observed a clear decrease in accessibility in BCG-vaccinated samples at Day 4 (Fig. 7E). Given the importance of NFKB1 as a defining transcription factor for the MyNFKB1^hi^ cell state, we examined accessibility of the *NFKB1* locus to determine if epigenetic regulation of *NFKB1* might play a role in its decreased expression in BCG-experienced myeloid cells and the downregulation of MyNFKB1^hi^ inflammatory genes. We found that multiple regions within the gene body decreased in accessibility with BCG in myeloid cells and remained lower than controls until at least 3 months post-vaccination (Fig. 7F). Overall, the single-nuclei data indicated a transient decrease in inflammatory MyNFKB1^hi^ phenotypes at 4 days post-BCG vaccination, with partial anti-inflammatory effects persisting up to 3 months.

Finally, to summarize our findings in both progenitor cells and myeloid progeny, we visualized chromVAR motif accessibility and gene expression scores of key TFs and gene modules presented in this work (Fig. 7G). In our data, control HSC were characterized by an oxidative phosphorylation and interferon-response expression program that peaked at Month 13, alongside a gradual increase in NRF1 motif accessibility over time. Upon the introduction of BCG, NRF1-motif containing regions became accessible early (at Day 4), and remained more open compared to control samples. Interferon response programs also peaked earlier (at Month 3) before falling to adult-like levels thereafter. In myeloid cells, we observed a distinct infant-specific cell state in control samples strongly associated with NF-κB activity and activation of the IL-1β inflammasome pathway. This corresponded with high NFKB1 motif accessibility up to Month 3, with high expression of the NF-κB regulated gene program in myeloid cells. These signatures followed similar patterns in HSC, suggesting shared regulation between progenitors and progeny cells. With BCG vaccination, NF-κB activity and target gene expression were attenuated in myeloid cells, especially at the Day 4 time point.

## Discussion

In this work, we present the first in-depth timeline of HSC development in human infants across the first year of life, including how they are affected by a potent, but safe, immune stimulus. While recent technological advances have enabled systems-immunology analyses in human infants (*13, 35, 36*), studies examining infant hematopoiesis have either consisted of few samples over a large age range or used cord blood as a source of HSC (*53, 83, 84*). Our data support earlier findings that gene expression in cord blood is not similar to that of neonatal PBMC (*13, 53*). Focusing on control neonatal samples, our results describe consistent trends in hematopoietic fate propensities and gene expression across infants. We observed an increased proportion of primitive HSC in early timepoints compared to adults, accompanied by high expression of genes associated with stemness and quiescence. Conversely, gene programs associated with differentiation, interferon response, and oxidative phosphorylation increased up to 13 months of age. These findings differ from studies of early-life hematopoiesis in mice, in which most HSC are cycling during the weeks after birth (*85, 86*), but concur with Zhang et al.’s 2022 study of human infant bone marrow, in which upwards of 75% of neonatal HSC were found to be in G0 phase of the cell cycle (*53*). Given the lack of existing resources for the study of human early-life HSC, and the lack of concordance with murine models, the HSC gene expression trends described in this work provide a valuable resource for further study of how age affects HSC—to define both the causes (e.g. environmental and microbial exposures, vaccination, allergens) and consequences (e.g. response to infections, immune memory, reconstitution potential) of transcriptional changes observed over early life.

Analysis of HSPC subpopulations revealed a unique circulating B-cell progenitor (BP) in neonatal samples and cord blood. There are two possible interpretations for this population and its relevance to immune development in early life. First, these cells could represent a wave of B-1 cell progenitors in neonates that subsides shortly after birth. B-1 cells are innate-like B cells best profiled in mice, with less well-described function in humans, that are particularly abundant in body cavities (*44, 87*). The majority of B-1 cells are produced in early life: murine fetal liver HSC can differentiate into B-1 cells, whereas adult bone marrow HSC cannot (*45, 88*). Given that mature B-1 cells are not found in appreciable numbers in the fetal liver (*89*), it has been hypothesized that B-1 cell progenitors migrate from the fetal liver to body cavities in the early weeks following birth; this timing coincides remarkably well with the peak abundance of circulating BP population observed in our data. A second possible interpretation relates to overall B cell abundance. B cells as a whole are highly abundant in infants, and increase in number over early life (*13, 40*). Maintaining this population likely requires high levels of B-lymphopoiesis, which would increase the likelihood of capturing B cell progenitors as a rare cell type in data from infant timepoints. Future work should determine if there is a relationship between these early B progenitor cells we describe and acute lymphoblastic leukemia, especially of the B cell precursor subtype, which is the most common form of leukemia in children (*90*). Unfortunately, the rarity of BP cells in our data does not allow for conclusive analysis of their cell fate. Future efforts to isolate and deeply characterize this cell population will be essential to determine its contribution to human B-cell ontogeny and disease.

In descendant myeloid cells, we were surprised to observe a dominant NF-κB-associated inflammatory phenotype (MyNFKB1^hi^) in both epigenetic and transcriptional data up to month 3 of life. Literature on myeloid cells in human infants is conflicting, with little consistency in reported cytokine responses to stimuli (*91*); of course, many of these studies rely on cord blood in place of newborn samples. We believe that, based on baseline intracellular cytokine data and correlation with whole-blood cytokine production capacity upon stimulation, the MyNFKB1^hi^ cells observed in our data have a greater capacity for TNF, IL-6 and IL-1β production than conventional myeloid cells. An abundance of myeloid cells primed for inflammatory cytokine production could offer an explanation for the increased susceptibility of young infants to hyperinflammatory complications during sepsis—a condition with a mortality rate between 11% and 19% in neonates (*92*). Despite the inconsistency of results in human samples, neonatal mice have been shown to produce much higher levels of myeloid inflammatory cytokines upon LPS or other TLR stimulation than adult mice (*93*). Consequently, when injected with an adjusted LPS dose, neonatal mice died within 24-48 hours of challenge, while adult mice survived (*93*). While cytokines like TNF, IL-1β, and IL-6 are necessary for the activation of host defenses against bacterial pathogens, dysregulated and high expression of these signaling molecules has been associated with many of the most damaging consequences of sepsis (*94*). Evolutionarily, an abundance of myeloid pro-inflammatory cytokines could help infants control early-stage bacterial infections in the absence of mature memory B and T cell responses. However, if an infection escapes immune control, such “all or nothing” neonatal innate immune response may contribute to inflammatory pathology and sepsis.

Having established a timeline of infant immune development in HSPC and the innate immune system, we then examined how BCG vaccination changes its course. First, BCG induced early IFN-γ signaling. The concept that BCG induces a Th1/IFN-γ cytokine response in the months following vaccination is well established in human adults and in mice, with multiple studies reporting that the heterologous protective effects of BCG are dependent on IFN-γ (*25, 28, 95*). The interferon response signature we observed in infant HSC at 3 months post-vaccination is similar to that observed in adults post-BCG immunotherapy, though not identical (*28*); these differences may derive from the route of administration (intravesical vs. intradermal) as well as age. There is also evidence in additional studies of vaccinated infant cohorts that early introduction of BCG induces a robust Th1 response that peaks 6-10 weeks after vaccination (*96–98*). Unexpectedly, this same signature appears in control samples at 13 months of age, implying that a wave of IFN-γ signaling in infancy is a consistent, and potentially developmentally programmed, phase of normal hematopoietic development. Notably, many Danish children start to attend daycare at one year of age (*99*), just before our final month 13 timepoint. This wave, which may coincide with a rise in antigen experienced lymphocyte populations over time, could represent a defining signal directing maturation of hematopoiesis by the end of the first year of life.

Conversely, in myeloid cells, we observed a transient reduction in myeloid inflammation shortly after BCG vaccination. While BCG vaccination is best known for protection from tuberculosis and the induction of innate immune memory (‘*trained immunity*’), it has also been shown to reduce systemic inflammation (*81, 82*). We see this effect in our data as a decrease in MyNFKB1^hi^ abundance 4 days post-vaccination, illustrated in both sequencing and flow cytometric assays. Changes in myeloid cell state at this early timepoint are especially interesting in light of clinical studies showing that BCG’s greatest protective effect is in the early neonatal period, primarily derived from a reduction in the mortality due to fatal neonatal sepsis (*100, 101*). This observation is supported by experiments in mice that demonstrate robust protection from polymicrobial sepsis at early timepoints following neonatal BCG vaccination (*102*). Our work suggests that myeloid cell state plays a role in this protection. The finding that BCG vaccination rebalances the excess production of sepsis-associated cytokines such as TNF, IL-1β, and IL-6 would support a model in which decreased inflammatory signaling may contribute to the reduced sepsis mortality post vaccination in newborns.

Overall, our data show that the protective effects of BCG against infant mortality may arise from multiple (not necessarily linked) mechanisms acting on HSC and immune progenitors. First, a short-term reduction in myeloid inflammatory transcription may decrease the risk of excessive systemic inflammation during infection in neonates. Second, the acceleration of an IFN-γ program induces HSC phenotypes that could reduce susceptibility of infants to infection. Our study offers a methodology and conceptual framework for future studies to chart the coordinated development of the infant immune system and hematopoiesis. With BCG as an example, this work highlights the potential of early exposure to controlled immunological stimuli to shape hematopoietic and immune programs in ways that may reduce the toll of infections, allergies, and other immune-mediated pathologies in children. Future work should further assess this model, the environmental factors that contribute to the reduction in NF-κB programs and cumulative IFN-γ signature in variable infant environments (e.g., industrial and rural), and the correlation between these signatures and resistance to infection and sepsis and other features of early life immunity.

## Funding

National Cancer Institute (NCI) award U54-CA274492 (D.P.)

Cancer Center Support grant P30-CA0008748 (D.P.)

Howard Hughes Medical Institute investigator (D.P.)

European Research Council (ERC) Advanced Grant 833247 (M.G.N.)

Spinoza Grant of the Netherlands Organization for Scientific Research (M.G.N.)

National Institutes of Health grant R01AI148416 (S.Z.J.)

Hevolution Foundation grant HF-GRO-23-1199240-4 (S.Z.J.)

American Society of Hematology (S.Z.J.)

## Author contributions

Conceptualization: C.J., L.S.H., S.Z.J., M.G.N.; Methodology: L.P., C.J., L.S.H., D.P.; Investigation: C.J., L.S.H., V.L., A.M.P., A.W.D., N.M.B.W., T.N.N.; Data Curation: L.P., C.J., L.S.H., T.N.N.; Formal Analysis: L.P., L.S.H., C.K.B.; Visualization: L.P., C.J., L.S.H., C.K.B.; Funding Acquisition: S.Z.J., M.G.N., D.P., P.A., C.S.-B.; Project Administration: L.P., C.J., L.S.H.; Supervision: S.Z.J., M.G.N., D.P., R.E.N., P.A., C.S.-B.; Writing – original draft: L.P.; Writing – review & editing: L.P., C.J., L.S.H., C.K.B., N.M.B.W., V.L., A.M.P., T.N.N., A.W.D., P.A., C.S.-B., R.E.N., D.P., S.Z.J., M.G.N.

## Competing interests

D.P. reports equity interests and provision of services for Insitro, Inc. M.G.N. is a scientific founder of TTxD, Lemba, Salvina and Biotrip. S.Z.J. is a scientific founder of Epistemyx, Inc. All other authors declare no competing interests relevant to this manuscript.

## Data, code, and materials availability

## Materials

This study did not generate new, unique reagents.

## Data and code

Code written for data processing, analysis, and visualization in this study will be made publicly available in a Zenodo repository (https://doi.org/10.5281/zenodo.18635596) upon journal publication. Processed snRNA/snATAC data files will similarly be made publicly available in a Zenodo repository (https://doi.org/10.5281/zenodo.18625985) upon journal publication. In the interim, access to both repositories is available upon reasonable request to the corresponding authors. Due to confidentiality concerns, raw human sequencing data used for genotype demultiplexing, as well as raw snRNA/snATAC sequencing files, will not be publicly released, but will be available to investigators upon reasonable request to the corresponding authors and fulfillment of a research data use agreement.

## Supplementary Materials

Materials and Methods

Figs. S1 to S8

Tables S1 to S8

## Materials and Methods

### Experimental design and study participants

#### Calmette (BCG) cohort

This cohort consists of 4,262 infants born in Rigshospitalet, Hvidovre Hospital, and Kolding Hospital in Denmark from October 2012 to November 2015. Written consent was obtained from the parents of all participants. The trial was approved by the Danish Data Protection Board, the Committees on Biomedical Research Ethics, and the Danish Medicines Agency, and can be found under the trial registration number NCT01694108. Children included in the trial were recommended the standard Danish vaccine schedule of three doses of DiTeKiPol/Act-Hib and Prevenar 13 at 3, 5 and 12 months. Due to a period of no available DiTeKiPol/Act-Hib during the trial, a subset of children were administered Infanrixhexa instead (*34*). Infants included in this study followed a normal distribution of birth weights and gestational age (Fig. S1A,B) in both BCG and control groups. One infant selected for sequencing in the control group was categorized as “very preterm” birth (< 37 weeks gestation), but had a birth weight within the normal range. Demographic information and applied assays for each sample are summarized in Supplementary Table 1.

### Blood collection

At the hospital, blood samples were drawn from infants 4 days post-randomization. Follow-up samples were collected at 3- and 13-month physical examinations. Whole blood was collected in 2 mL Lithium Heparin tubes, BD Vacutainer (BD Biosciences #368494).

### Progenitor input enrichment sequencing

Cryopreserved PBMCs from 8 donors or cord blood mononuclear cells (CBMCs) from 4 donors were thawed in a 37 °C water bath and transferred into 15 mL conical tubes containing 10 mL of thawing medium (RPMI supplemented with 10% FBS at room temperature) per batch. After gentle inversion, 10–15% of cells from each sample were set aside for live cell sorting. All samples were centrifuged at 350 × g for 10 min at room temperature. The reserved 10–15% aliquots were resuspended in MACS buffer (1× PBS supplemented with 2 mM EDTA and 0.5% BSA) and kept on ice. The remaining cells from all donors in the same batch were pooled and enriched for CD34⁺ cells using the CD34 MicroBead Kit (Miltenyi #130-046-702) and MS Columns (Miltenyi #130-042-201) following the manufacturer’s protocol. Pooled cells were stained at 4°C for 30 min with antibodies against lineage markers (1:25, BV510, BioLegend #348807) and CD34 (1:100, FITC, Miltenyi #130-113-178) in the dark. After a wash with MACS buffer, live pooled lineage⁻ CD34⁺ (Lin^-^ CD34^+^) cells and individual PBMCs / CBMCs were resuspended in 7-AAD Viability Staining Solution (1:100, BioLegend #420404). Without additional washing, Lin⁻ CD34⁺ cells were sorted directly into individual PCR tubes on a FACSAria™ Fusion Flow Cytometer (BD Biosciences). Live PBMCs or CBMCs from the reserved aliquots were then sorted into the same tube to achieve a final PBMC / CBMC:Lin⁻ CD34⁺ ratio of 7:1 to 1:1. Nuclei were isolated using the low-input protocol from 10x Genomics (CG000365 | Rev C). Multiomic libraries were prepared to recover 4,000–15,000 nuclei per sample lane, using the Chromium Next GEM Single Cell Multiome ATAC + Gene Expression Reagent Bundle (10x Genomics #1000283) and the Chromium Next GEM Chip J Single Cell Kit (10x Genomics #1000234), according to the manufacturer’s user guide (10x Genomics, CG000338 | Rev F). cDNA and final libraries were quantified using the High Sensitivity DNA Kit (Agilent #4067-4626). Libraries were sequenced on an Illumina NovaSeq X Plus platform at the Genomics Resources Core Facility, Weill Cornell Medicine (New York, NY, USA).

### Low-pass whole genome sequencing

Low-pass whole genome sequencing (∼3 Gb, ∼1× coverage) was performed on ∼1 × 10⁶ cells from each PBMC and CBMC donor by Genewiz from Azenta Life Sciences (South Plainfield, NJ, USA). FASTQ files were delivered for single nucleotide polymorphism (SNP) identification. Briefly, sequencing reads were trimmed using TrimGalore! and aligned to the human reference genome (hg38) using bwa-mem2. Reads with low mapping quality were filtered, and duplicates were marked and removed using Picard MarkDuplicates. Variant calling was performed with bcftools on both original and downsampled BAM files, followed by normalization and filtering to generate variant call format (VCF) files.

### Whole-blood stimulation and ELISA

Whole blood samples were cultured in sterile round-bottomed 96-well plates in triplicate using culture medium (Roswell Park Memorial Institute (RPMI) 1640) supplemented with gentamicin 5 μg/mL, glutamax 10 mM and pyruvate 10 mM. 140 μl culture medium was added to 40 μl whole blood and 20 μl of stimuli to achieve a final concentration of: culture medium (negative control), LPS 100 ng/mL (Lipopolysaccharide from *Escherichia coli* 055:B5, L-4005, Sigma), PHA 100 μg/mL (phytohemagglutinin, L-1668, and L-8902, Sigma), HK *E. coli* 1 × 10^6^ microorganisms/mL (ATCC 25922), HK *Streptococcus pneumoniae* 1 × 10^6^ microorganisms/mL (ATCC 49619), and HK *Candida albicans* 1 × 10^6^ microorganisms/mL (MYA 35730). Samples were incubated at 37 °C and 5% CO_2_; cytokine measurements were performed after 24h for myeloid-derived cytokines (IL-6, IL-1β, and TNF-α) and 48h for lymphoid-derived cytokines (IFN-γ, IL-10, and IL-22).

Cytokines were measured in culture supernatant using commercially available ELISA kits from Sanquin (IL-6, IL-10 and IFN-γ) and R&D Systems (TNF-α, IL-1β, and IL-22) according to instructions of the manufacturer. The ELISA dynamic ranges were: IL-1β: 3.9–500 pg/ml, TNF-α: 7.8–1000 pg/ml, IL-6:1.6–200 pg/ml, IFN-γ: 3.9–500 pg/ml, IL-10: 2.3–300 pg/ml, and IL-22: 15.6–2000 pg/ml. Sample dilutions used were: IL-1β 10×, TNF-α 10×; IL-6 from RPMI 10×; IL-6 from all other stimuli 100×; IFN-γ from PHA stimulation 50×; IFN-γ from all other stimuli 2×; IL-10 2×, and IL-22 2×. Reported cytokine measurements are adjusted for dilution factor.

### Serum protein Olink

Inflammatory protein concentrations were measured using the Olink Target 96 Inflammatory panel platform at all three time points. Protein abundances were quantified using EDTA plasma samples. Immunoassays used by Olink are based on the Proximity Extension Assay (PEA) technology (*103*). This technology uses oligonucleotide-labelled antibodies binding to their respective target proteins in the samples. When two antibodies are brought into close proximity, a polymerase chain reaction (PCR) reporter sequence is formed by DNA polymerization, which is subsequently quantified by quantitative real-time polymerase chain reaction (qPCR). To avoid batch effects, the BCG and the control group samples were randomly distributed across plates. Each plate included interplate controls which are used to adjust any potential plate differences. The Olink data are reported as Normalized Protein eXpression (NPX) values on log 2 scale, a relative quantification unit as protein concentration.

### MyNFKB1^hi^ stimuli

RSV A2 and *S. pneumoniae* were kindly provided by Dr. R. Uijen and Dr. M. de Jonge and were cultured as described previously (*104, 105*). In brief, RSV A2 was cultured on 70% confluent HeLa cells (ATCC CCL-2) for 4 days. Cells were scraped and the suspension was centrifuged to remove cellular debris. RSV was purified by ultracentrifugation over a 30% sucrose layer for 90 min at 20,000g, snap frozen, and stored at -80 °C until use.

Virus titer was quantified by titration on HeLa cells. Near confluent HeLa cells (80–90%) were infected with ten-fold dilutions of RSV virus for 1 hour, after which cells were washed once with PBS and incubated for 72 hours at 37 °C, 5% CO_2_. RNA was harvested by lysing cells in 50µL iScript RT-qPCR sample preparation reagent (Bio-Rad). RNA was transcribed using the iScript cDNA synthesis kit (Bio-Rad), and qPCR was performed using the SsoAdvanced Universal SYBR Green Supermix (Bio-Rad). TCID50 (50% tissue culture infectious dose) was calculated based on the number of positive wells per dilution.

*S. pneumoniae* serotype 4 TIGR4 was grown and prepared as previously described (*106, 107*). *S. pneumoniae* was inactivated using β-propiolactone (BPL) by adding 0.025% BPL to the bacterial stock followed by incubation for 16 hours at 4°C under continuous slow shaking. Next, the stock was incubated for 4 hours at 37 °C to hydrolyze the BPL and stored at -80 °C until further use.

### MyNFKB1^hi^ flow cytometry

Frozen PBMCs were thawed in a 37 °C water bath and transferred into conical 15mL tubes filled with pre-warmed RPMI supplemented with 2mM GlutaMAX, 1mM pyruvate, 100U/mL penicillin-streptomycin, and 20% FCS, hereafter referred to as RPMI+. After centrifugation, cells were washed twice more and resuspended in RPMI+ for counting. PBMCs at a concentration of 5×10^6 cells/mL were dispensed into a round-bottomed 96-well polypropylene plate (VWR, cat# 736-0117) and stimulated with LPS 10ng/mL, R848 10μg/mL, *S. pneumoniae* or live RSV A2 at a multiplicity of infection (MOI) of 10. Cells were cultured at 37 °C in a humidified atmosphere of 5% CO_2_ for 4 hours in the presence of a Golgi plug (Brefeldin A).

Cells were transferred to v-bottom plates and washed with PBS. Cells were incubated with ViaKrome viability dye for 5 min at RT, then washed with PBS + 1% bovine serum albumin (BSA). Surface staining of CD44 and CD83 was performed prior to fixation and permeabilization of the cells, followed by subsequent staining for intracellular cytokine and other markers of interest (antibodies used are listed in the reagent table). Samples were analyzed on a CytoFLEX LX flow cytometer (Beckman Coulter).

### Quantification and statistical analysis

#### Whole-blood stimulation analysis

Cytokine stimulation data was analyzed using R version 4.4.1. Protein concentration measurements by ELISA were log-transformed to approximate a normal distribution. Analysis was restricted to the 66 samples with matched measurements across all three control timepoints. For each cytokine of interest, comparisons between timepoints were performed using ANOVA followed by paired post-hoc comparisons between timepoints with Holm correction for multiple testing.

To assess the relationship between myeloid cell proportions in the scMultiome data and cytokine concentrations, we conducted a non-parametric correlation analysis at each study timepoint (Day 4, Month 3, and Month 13). Spearman’s rank correlation coefficients (ρ) were calculated for each timepoint, correlating myeloid cell state frequencies with log-transformed cytokine levels. Data visualization was performed using the *corrplot* package in R (*108*).

#### Serum protein analysis

Serum protein data was analyzed using R version 4.4.1. Normalized Protein Expression (NPX) values from Olink proteomics for proteins of interest in n=128 control and n=178 BCG infants were compared between BCG and control conditions for each timepoint using a two-sample Student’s t-test followed by Holm multiple testing correction.

#### MyNFKB1^hi^ flow cytometry

All flow cytometry data were analyzed using FlowJo software (version 10.10.0). Each file was gated on according to the strategy shown in Supplemental Figure 8. Graphs were made using GraphPad Prism 10.

#### Alignment and demultiplexing of single-nuclei Multiome sequenced data

All sequencing lanes were preprocessed with the Cell Ranger ARC 2.0.2 pipeline using the GRCh38 reference genome (sample demultiplexing, barcode processing, alignment of reads, and counting of transcripts). Initial cell filtering of the raw feature-barcode matrix was performed using EmptyDrops Multiome (*109*). For genotype demultiplexing, aligned gene expression .bam files from lanes containing the same set of genotypes were combined using mergeBams (*110*), keeping only barcodes that passed initial EmptyDrops filtering. Souporcell was run on each merged .bam with default parameters (*111*), using the expected cell barcodes and number of genotypes within each merged .bam as additional input. Souporcell predicted genotypes were matched to known genotypes based on the maximum number of shared SNPs.

Filtered GEX count matrices were then loaded into Python using Scanpy 1.10.4, and additional cell filtering was performed per sequencing lane by manual inspection of embeddings of gene expression data (standard log and median normalization, PCA, UMAP, and PhenoGraph clustering) (*112*). Alongside standard Scanpy QC metrics, Scrublet was run on raw counts from each sequencing lane for doublet detection (*113*). Poor-quality cell clusters were removed based on examination of total GEX counts per cell, Scrublet doublet score, and Souporcell assignment (Souporcell-predicted doublets or unassigned cells). All Souporcell-predicted doublets or unassigned cells were then removed.

#### CellTypist annotations

Cell type annotations of the adult control samples included in this dataset were used to guide annotation of infant and cord blood cells using CellTypist (*114, 115*). The adult data was downsampled to 9,980 cells to use as a training set, including higher proportions of rare cell types than present in the full adult data by setting the balance_cell_type parameter in CellTypist’s downsample_adata function to True. The remaining 26,737 adult cells were withheld as a testing set. Raw RNA count data was rescaled to 10,000 counts per cell using scanpy.pp.normalize_total and then log-normalized with a pseudocount of 1. A CellTypist model was trained to predict cell type from normalized counts using default parameters, and achieved 0.96 accuracy, 0.84 precision, and 0.93 recall against the testing set. The trained model was then applied to scaled and log-normalized GEX counts of the remaining snMultiome data to generate cell type predictions for cells from all age groups.

#### MrVI model training

To address the challenge of generating fine-grained cell type annotations consistent across age groups, without ignoring real age-group associated variation in the data, we chose to train a MrVI model on our single-nuclei GEX data (*37*). MrVI is a deep-generative model that maps cells into two latent spaces: a u-space designed to capture cell-state effects, and a z-space that captures effects of target covariates such as biological sample and condition. Both latent spaces should remain unaffected by nuisance covariates such as batch, which are specified using the ‘batch’ key in model training.

Raw GEX counts of filtered barcodes from all samples were concatenated into a full-dataset cell count matrix. 2,000 highly-variable genes (HVG) were selected with the ‘seurat_v3’ flavor of scanpy.pp.highly_variable_genes using the concatenation of condition and timepoint as the batch key. Mitochondrial and ribosomal genes were removed from consideration for HVG selection. Using biological sample as the sample key, and the concatenation of sequencing lane and sex as the batch key, MrVI models were trained on this count data across a range of possible u-space and z-space dimensionality: selecting from u = (10,20,30) and z = (10,20,30,50,100) dimensional latent spaces where z >= u.

To evaluate robustness of cell type annotations across models, PhenoGraph (k = 30) was run on each model’s u-space. Cells were annotated as the majority CellTypist predicted label within each cluster. Then, cell annotations were compared across models using Adjusted Rand Index (ARI) to quantify the similarity of cell labels between runs. Final annotations and UMAP visualization of the u-latent used in this paper were calculated from the u=20 z=30 MrVI model.

#### Single-cell multiome ATAC preprocessing

Single-cell ATAC data was analyzed using the Pe’er lab v1.0.1 fork of ArchR (*116*). ATAC fragment files from CellRanger ARC output and filtered barcodes were used to generate arrow files and a corresponding ArchR project. ArchR’s iterative LSI implementation was run on a genome-wide matrix of 500bp tiles to generate a 29-dimensional SVD embedding for UMAP visualization. Total dataset cell type annotations were used as the groupBy parameter for ArchR pseudo-bulk replicates and peak calling. Peaks were annotated to motifs using the CISBP database. ChromVAR motif enrichment score per cell was determined using ArchR’s addBgdPeaks and addDeviationsMatrix functions. Marker peaks were found for clusters of interest and evaluated for motif enrichment using the getMarkerFeatures and peakAnnoEnrichment functions as described in ArchR documentation.

#### Milo differential abundance

The differential abundance method Milo was used to test for enrichment of infant vs. adult cells across all cell types (*38*). Milo neighborhoods were generated (prop = 0.1, k = 30, d = 20) using the MrVI u-latent space to best capture cell-type information. Differential abundance testing was conducted between all control infant samples vs. all control adult samples, producing a logFC value and false discovery rate (FDR) per neighborhood. Each neighborhood was annotated to a cell type using Milo’s annotateNhoods() for visualization of neighborhood logFC distributions across cell types.

#### Subsetting and reprocessing of HSPC scMultiome data

The 38,415 cells labeled as HSPC in the final MrVI model were subset from the dataset for in-depth analysis. These HSPC were subjected to finer cell filtering to enable later pseudotime analysis, given that constructing cell-fate trajectories can be quite sensitive to cell quality. Any cell with GEX counts less than 1000 or greater than 30,000 were removed from the data subset, leaving 36,417 remaining HSPC.

In order to emphasize cell-state associated genes, which should be variable within all age groups, over age-associated genes, which should vary across age-groups, we took a different approach to HVG selection in the HSPC subset. 3,000 total HVG were calculated from the HSPC GEX raw count data using the ‘seurat_v3’ flavor of scanpy.pp.highly_variable_genes with age group as the batch key. Ribosomal and mitochondrial genes were excluded from consideration. This generates a ‘highly_variable_nbatches’ column in the AnnData var slot, indicating the number of age groups that a gene is found to be highly variable in. We kept the 1,090 HVG found to be variable in all three (cord blood, infant, and adult) age groups for use in downstream analysis.

Although MrVI worked well for age group integration in the full dataset, the mixture of Gaussians prior it uses to generate a cell-state driven u-space is better suited to discrete cell type clusters than smooth cell fate trajectories. Therefore, we chose to instead use Scanorama to integrate age groups within HSPC GEX data (*117*). After standard log and median normalization, the HSPC GEX data was split by age group and then integrated using scanorama.integrate_scanpy with dimred=30 and alpha=0.01. This generated a 30-dimensional ‘Scanorama’ embedding of the HSPC GEX data, which we used to generate the neighbor graph and visualize with a force-directed layout. For HSPC subtype annotations, PhenoGraph clusters (k=30) were generated from the Scanorama embedding and manually labeled according to expression of canonical marker genes and mean chromVAR motif accessibility score per cluster.

After characterization of the B-progenitor cluster, these cells were removed to enable finer characterization of major trajectories. The remaining 36,083 HSPC were reprocessed with the same methods and parameters as the full HSPC object.

#### HSPC pseudotime analysis

HSPC cell fate trajectories were inferred using Palantir (*3*). Diffusion maps were generated from the HSPC Scanorama embedding, and the top 7 eigenvectors were selected for use in trajectory analysis based on manual inspection of eigenvalues. Start and end cells were manually selected based on expression of key HSPC marker genes, with *HLF* gene expression characterizing the HSC starting cell, and LMPP, GMP, ERP, and BEM endpoints defined by *DNTT*, *MPO*, *GATA1*, and *HDC* expression respectively. After running palantir, trajectory branch masks were generated using palantir.presults.select_branch_cells (q=.005, eps=.01) to visualize cell densities, gene expression, and motif accessibility along distinct lineage pseudotime. To combat dropout in gene expression visualizations, MAGIC (t = 3) was applied to the log-normalized count data and Scanorama embedding (*118*). MAGIC imputed data was used only for visualization purposes, and is described in the main text and figure legends when applied.

#### Multiome snRNA differential gene expression

Differential gene expression in the scMultiome data was conducted using MAST, a hurdle model that accounts for the many zero-counts in single-cell/nuclei data (*119*). For each comparison, we used MAST to model log-normalized snRNA count data as a function of condition and cellular detection rate (the fraction of genes detectably expressed in each cell). In initial analysis, we noticed that variation in the number of female and male cells from different conditions resulted in sex-linked genes such as *XIST* being ranked as significant differential genes. Therefore, we added sex as an additional covariate to the model. We used a likelihood ratio test to determine differential expression between conditions, returning a false discovery rate (FDR) and natural log-fold change per gene, which was converted to log_2_ fold-change (log_2_FC) by dividing by the natural log of 2.

Given the known risk of false positives in single-cell/nuclei differential expression we used relatively strict cutoffs of FDR < 0.001 and absolute log_2_FC > 0.1 to define genes as significant in each comparison (*120*). Significantly up or down-regulated genes were compared to GO Biological Process 2021 genesets using EnrichR to aid in biological interpretation (*121*). For specific comparisons to genesets in literature, GSEA was run on differential expression results ranked according to sign(log_2_FC) * -log_10_(FDR) (*122*).

#### HSC differential gene clustering

Differential genes in HSC were clustered according to their change in expression over control timepoints. First, as a form of feature selection, only genes found to be differentially expressed between at least one infant control timepoint vs. adult were included in this analysis: a total of 3,484 genes. Gene log_2_FC values from the comparison of each timepoint to adult (with adult vs. adult log_2_FC set to 0) were z-scored across timepoints; this focused clustering results on the pattern of expression over time, rather than magnitude of change. These z-scores were clustered with k-means to produce 7 gene clusters, which were ranked and labeled 0-6 according to cluster size.

To produce cluster expression scores per individual, cells were scored using scanpy.tl.score_genes according to each cluster geneset, and then the mean cell score in HSC was calculated per individual and timepoint. Samples with <= 20 HSC were excluded from visualization. For gene cluster interpretation, cluster member genes were compared to GO Biological Process 2021 genesets using EnrichR (*121*).

#### PCA analysis of HSC transcription

Cells annotated to HSC were subset from the overall HSPC transcriptional data object. Mitochondrial and ribosomal genes, as well as genes expressed in fewer than 10 HSC, were excluded from this analysis. For feature selection, the top 5,000 highly variable genes were determined using scanpy.pp.highly_variable_genes (seurat_v3 flavor, run using raw counts). Samples were pseudobulked according to mean-expression of log-normalized counts for these 5,000 highly variable genes. Pseudobulks with <= 20 cells were excluded. PCA was run on this sample-gene matrix and the top two PCs were visualized to describe overall variation in HSC gene expression.

#### Subsetting and reprocessing of myeloid multiome data

All cells annotated as CD14 Monocytes, CD16 Monocytes, or cDC were isolated from the total dataset. ArchR’s iterative LSI implementation was rerun on this subset with the same parameters previously described, and the resulting 29 SVD dimensions were visualized using UMAP. Myeloid clusters were generated by running PhenoGraph (clustering_algo = ‘leiden’, k=30) on this embedding. Given the embedding clearly captured cell state, ATAC SVD neighbors were used for MAGIC (t=3) imputation of expression data. Diffusion maps were calculated from the ATAC SVD embedding to identify major axes of epigenetic variation in myeloid cells (*76*). Diffusion Component 1 associated genes were found by ranking genes according to their Pearson correlation coefficient between Diffusion Component 1 and log-normalized gene expression vectors. Myeloid cells were categorized as MyNFKB1^hi^ or MyConv based on the mean Diffusion Component 1 value across all cells in a cluster: if the mean value exceeded 0.005, the cluster was labeled as MyNFKB1^hi^.

In initial data exploration, we saw a clear influence of inflammatory gene expression in both RNA and ATAC embeddings of myeloid cells. We chose to focus on the ATAC embedding, given the clear NFKB-driven epigenetic regulation of myeloid cells in this data. However, we also included an RNA visualization in supplementary figures to underscore the consistency of the myeloid inflammatory phenotype across data types. To generate this, 2,000 HVG were selected across myeloid cells using scanpy.pp.highly_variable_genes (seurat_v3 flavor, run using raw counts, excluding mitochondrial and ribosomal genes), and PCA was run on log-normalized RNA data in myeloid cells. Given that the top 165 PCs captured 60% of the variance, these PCs were used to generate a neighbor graph (k=30) and visualized with UMAP.

#### In-silico validation of MyNFKB1^hi^ signature

To determine if the MyNFKB1^hi^ myeloid gene signature existed in external infant datasets, we downloaded raw scRNA count matrices from a dataset of PBMC collected from two-month-old infants pre- and post-routine vaccinations (*80*). We included only the pre-vaccination timepoints in our analysis so as not to confound results with the possible effects of vaccination. Count matrices were loaded into Python using Scanpy 1.10.4. For cell filtering, cells with less than 10^3.5^ or greater than 10^4.25^ counts per cell were removed, alongside cells with more than 20% mitochondrial content. DoubletDetection was run on raw counts from each sample with standard parameters, and predicted doublets were removed (*123*).

To provide a comparison to adult cells, we also loaded in the healthy donor 10k PBMC dataset from 10x Genomics (*124*). For additional cell filtering, cells with less than 10^3.6^ or greater than 10^4.5^ counts per cell were removed, alongside cells with more than 15% mitochondrial content. DoubletDetection was run on raw counts and predicted doublets were removed.

To ensure the fairest “apples to apples” comparison of expression levels, we downsampled counts from both datasets to a goal of 6,000 mean counts/cell. We then combined the datasets by concatenating raw count matrices from all samples; this left us with 33,932 cells and 31,915 genes for joint analysis. 2,000 HVG were found using the seurat_v3 flavor of scanpy.pp.highly_variable_genes with age group as the batch key. Mitochondrial and ribosomal genes were removed from being candidate HVG. Counts were normalized using standard log and median normalization. PCA was run with 500 total PCs, and the top 53 PCs (explaining 60% of the variance) were kept for downstream calculations. To identify myeloid cells, PhenoGraph was run with k=30, and *LYZ*^+^ clusters were annotated as myeloid. These myeloid cells were scored using scanpy.tl.score_genes against the top 100 MyNFKB1^hi^ marker genes found by MAST differential expression analysis of MyNFKB1^hi^ vs. MyConv cells in the BCG infant dataset.

#### Pseudo-bulk ATAC analysis

For HSC / MPP and myeloid cells respectively, pseudo-bulk ATAC counts per sample were exported from the ArchR project using getGroupSE(), removing pseudo-bulks with fewer than 40 cells. For each cell type, peaks with no samples having greater than 50 reads, and peaks with less than 300 total reads across samples, were removed, leaving an accessible background set of 111,673 HSC peaks and 51,143 myeloid peaks. Peaks were annotated to nearest genes using ChIPseeker (*125*). The sample-peak matrix per cell type was analyzed using the DESeq2 differential accessibility pipeline (*126*), comparing all control infant timepoints to adult and comparing BCG to control. For identifying motifs within individual peaks (i.e. counting of NRF1 motifs within the Day 4 BCG vs. Control differential peak analysis in HSC), existing ArchR peak-motif annotations were used.

#### HOMER Motif Enrichment

To link Figure 3 HSC gene clusters to peaks, accessible peaks in pseudobulk HSC were filtered according to proximity to cluster genes. Specifically, a cluster’s set of associated peaks was generated by subsetting accessible HSC background peaks to peaks annotated by ChIPSeeker as proximal to any gene in that cluster. Peaks annotated as distal intergenic were excluded. Some cluster genes had no associated peaks in this analysis: 34 genes from Cluster 3, and 12 genes from Cluster 6. Motif enrichment analysis was then performed on each cluster’s associated peaks using HOMER findMotifsGenome.pl, which effectively scans the target (cluster-associated peaks) and background (HSC accessible peaks) sequences against a library of reliable motifs, returning motifs enriched in the target set with a p-value less than 0.05 (*127*).

HOMER was also used for motif enrichment analysis of significant differential peaks generated from pseudobulk-ATAC DESeq2 analysis. In this case, for each comparison, genomic coordinates of the set of significant (FDR < 0.05) peaks from DESeq2 output were exported and compared to the background set of accessible peaks in that cell type using HOMER findMotifsGenome.pl.

---

**Fig. S1.**
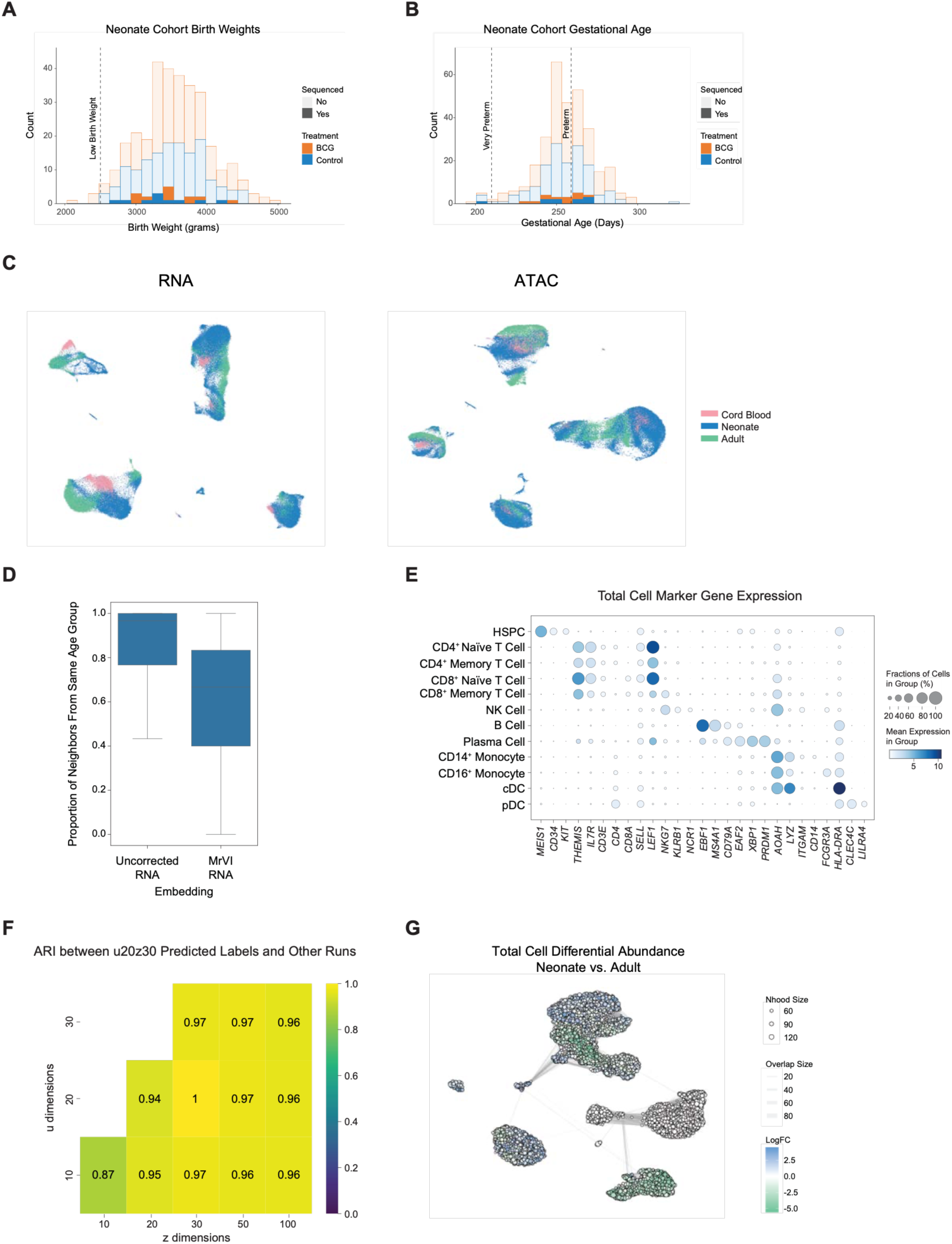
Heterogeneity in age group immune profiles. (**A**) Histogram of birth weight for infants included in the Calmette cytokine stimulation cohort, colored by treatment. Opacity indicates infants included in the PIE-seq sequencing dataset. Dotted line indicates the standard definition of low birth weight (<2500g). (**B**) Histogram of gestational age for infants included in the Calmette cytokine stimulation cohort, colored by treatment. Opacity indicates infants included in the PIE-seq sequencing dataset. Dotted lines indicate the standard definitions of preterm (<37 weeks gestation) and very preterm (<30 weeks gestation) birth. (**C**) UMAP visualizations of snRNA and snATAC PIE-seq data prior to sample integration, colored by age group. (**D**) Box plot of the proportion of each cell’s nearest neighbors in the snRNA embedding visualized in (C) that belong to the same age group, compared to nearest neighbors in the MrVI *u*-space. (**E**) Dot plot of marker gene expression (columns) by cell type annotation (rows) for the full PIE-seq dataset. Dot size represents the number of cells per group expressing each gene, and dot color represents log-normalized expression level. (**F**) Heatmap of adjusted rand-index (ARI) for cell type annotations between the MrVI model selected for final cell type annotations (*u* = 20, *z* = 30) and MrVI models trained with varying *u* and *z* dimension parameters, with rows indicating the number of *u* dimensions and columns indicating the number of *z* dimensions. (**G**) Milo results displayed on integrated snRNA UMAP as in Fig. 1C. Dots represent Milo neighborhoods, with dot size indicating the number of cells per neighborhood, and dot color indicating neighborhood logFC comparing all control infant to adult samples. Width of lines connecting neighborhoods represent the number of shared cells between neighborhoods. Positive LogFC indicates local enrichment of infant cells. LogFC of HSPC neighborhoods has been excluded, given that PIE-seq artificially enriches for HSPC.

**Fig. S2.**
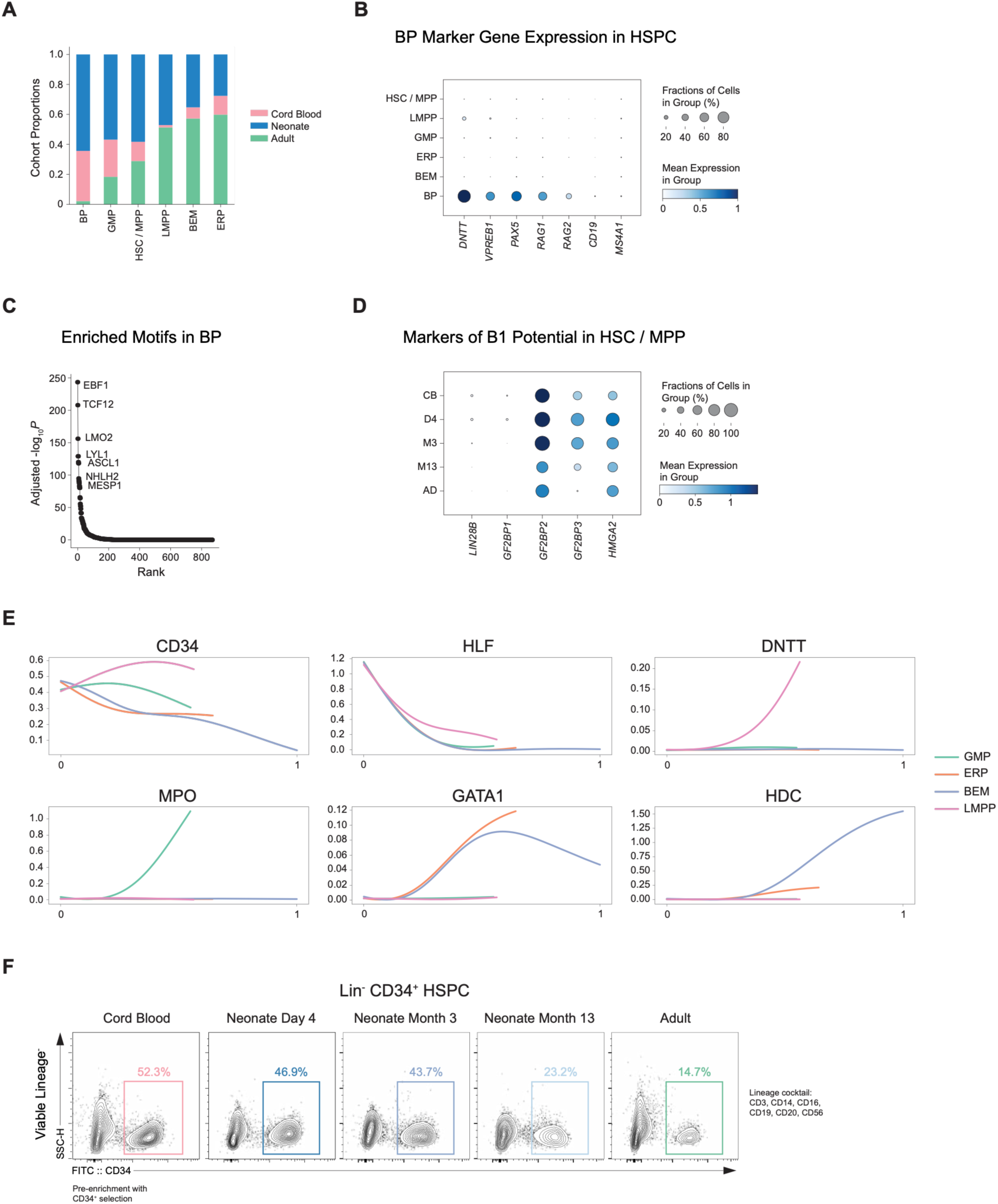
Differentiation potential of neonatal HSPC. (**A**) Bar plot of age group proportions within each HSPC subtype cluster. (**B**) Dot plot of marker gene expression (columns) by cell type annotation (rows) for selected BP marker genes. Dot size represents the number of cells per group expressing each gene, and dot color represents log-normalized expression level. (**C**) Rank plot of motif enrichment within snATAC marker peaks for BP compared to all other HSPC, with -log_10_P-adjusted on the y-axis and rank on the x-axis for each motif. (**D**) Dot plot of expression of markers of B-1 potential (columns) in HSC / MPP cells across control timepoints (rows). Dot size represents the number of cells per group expressing each gene, and dot color represents log-normalized expression level. (**E**) Palantir imputed gene trends over pseudotime for hematopoietic marker genes. (**F**) Flow cytometry plots of PIE-Seq Lin^-^CD34^+^ HSPC sorting from control samples across all time points. Boxes indicate the proportion of CD34^+^ cells within the Lin^-^compartment following pre-enrichment with CD34^+^ selection. *n* = 4-8 samples pooled per timepoint.

**Fig. S3.**
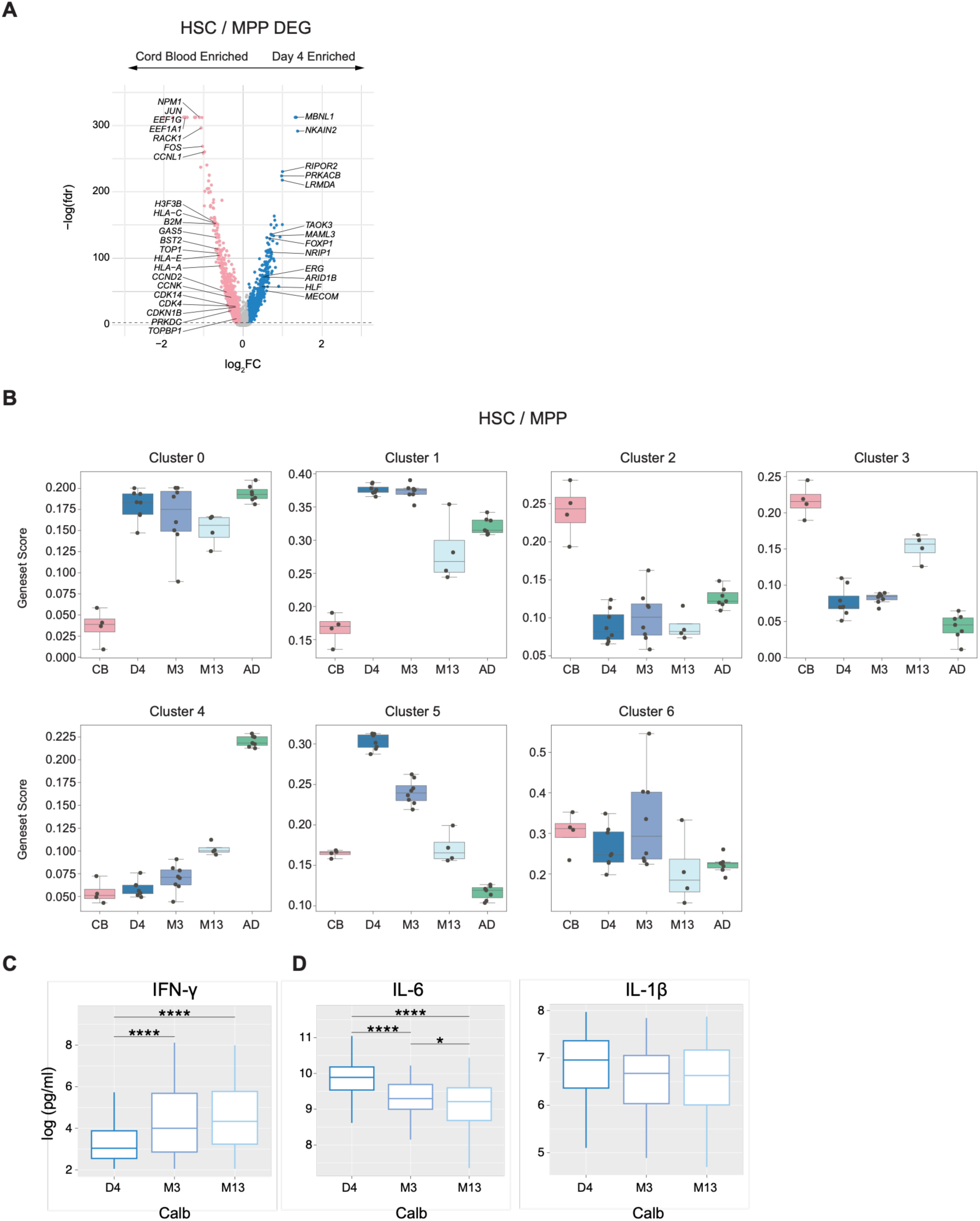
HSC gene expression programs across timepoints. (**A**) Differential gene expression results in HSC/MPP comparing Day 4 to cord blood samples. Volcano plot shows log_2_FC and - log(FDR) per gene, with genes with abs(log_2_FC) > 0.1 and FDR < 0.001 marked as significant. (**B**) Box plots of per sample expression score in HSC/MPP of Figure 3 gene clusters across control timepoints. (**C**) Log-normalized concentration (pg/mL) of IFN-γ upon 96h whole-blood stimulation with *C. albicans* of blood samples from n=66 control infant samples collected at 4 days, 3 months, and 13 months post-randomization. Statistical significance was assessed using a repeated-measures one-way ANOVA followed by paired post-hoc comparisons between timepoints with Holm correction for multiple testing. p > 0.05 = ns, ∗p ≤ 0.05, ∗∗p ≤ 0.01, ∗∗∗p ≤ 0.001. (**D**) Log-normalized concentrations (pg/mL) of IL-6 and IL-1β upon 24h whole-blood stimulation with *C. albicans* of blood samples from n=66 control infant samples collected at 4 days, 3 months, and 13 months post-randomization. Statistical significance was assessed using a repeated-measures one-way ANOVA followed by paired post-hoc comparisons between timepoints with Holm correction for multiple testing. p > 0.05 = ns, ∗p ≤ 0.05, ∗∗p ≤ 0.01, ∗∗∗p ≤ 0.001.

**Fig. S4.**
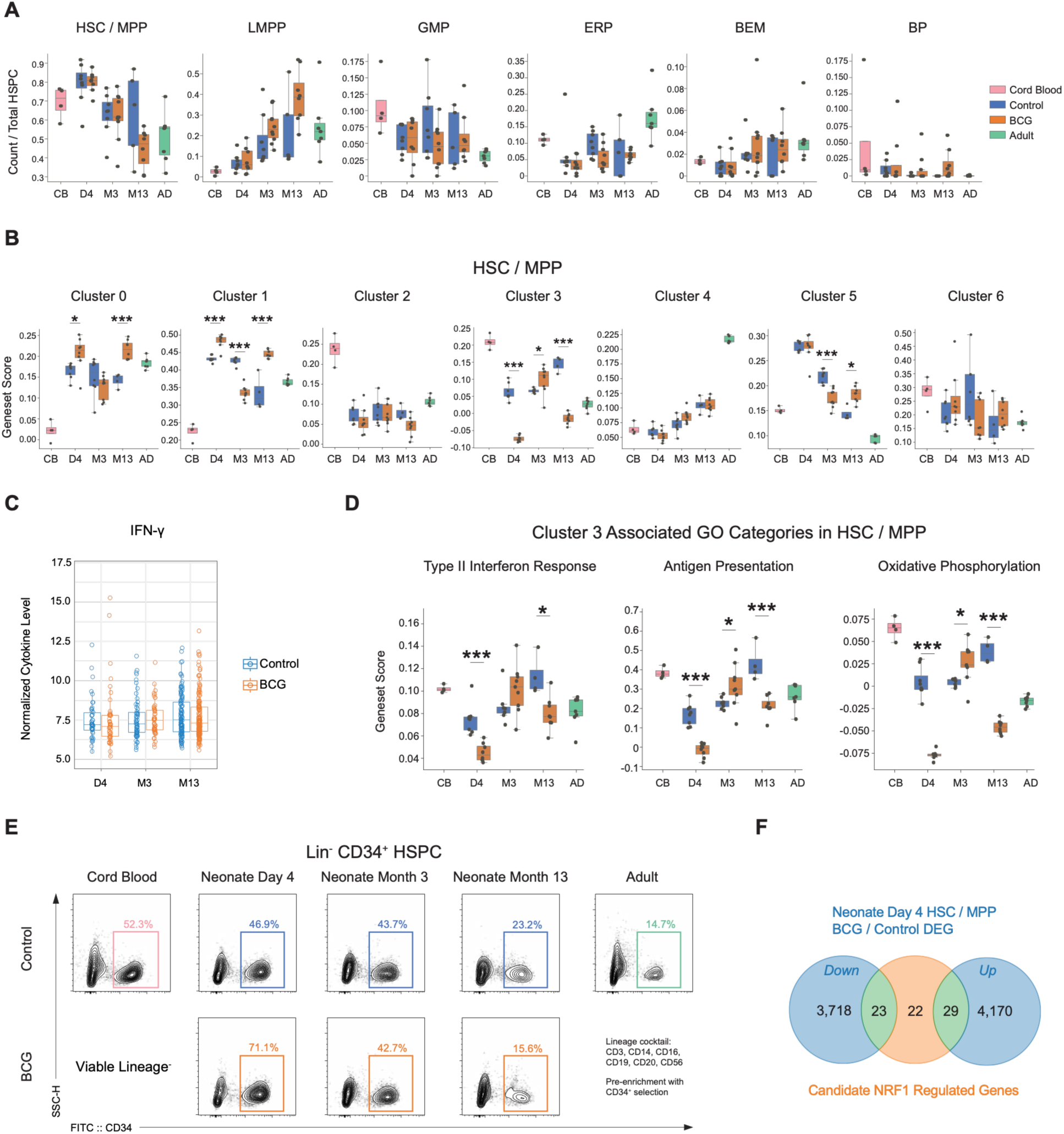
BCG vaccination-associated phenotypes in neonatal HSC. (**A**) Box plots of HSPC subtype proportion out of total HSPC for all samples, grouped by condition and timepoint. Differences between BCG and control groups at each timepoint were assessed using an unpaired two-sided Student’s *t*-test, with *p*-values adjusted for multiple comparisons using the Holm method. (**B**) Box plots of per sample expression score in HSC/MPP of Figure 3 gene clusters, grouped by condition and timepoint. Differences between BCG and control groups at each timepoint were assessed using an unpaired two-sided Student’s *t*-test, with *p*-values adjusted for multiple comparisons using the Holm method. p > 0.05 = ns, ∗p ≤ 0.05, ∗∗p ≤ 0.01, ∗∗∗p ≤ 0.001. (**C**) Box plot of Olink normalized protein expression level from serum of n = 128 control and n = 178 BCG-vaccinated infants at Day 4, Month 3, and Month 13 timepoints post-randomization. (**D**) Box plots of per sample expression score in HSC/MPP of Cluster 3 associated GO terms defined in Figure 3, grouped by condition and timepoint. Differences between BCG and control groups at each timepoint were assessed using an unpaired two-sided Student’s *t*-test, with *p*-values adjusted for multiple comparisons using the Holm method. p > 0.05 = ns, ∗p ≤ 0.05, ∗∗p ≤ 0.01, ∗∗∗p ≤ 0.001. (**E**) Flow cytometry plots of PIE-Seq Lin^-^ CD34^+^ HSPC sorting from control and BCG-vaccinated samples across all time points. Boxes indicate the proportion of CD34^+^ cells within the Lin^-^ compartment following pre-enrichment with CD34^+^ selection. *n* = 4-8 samples pooled per timepoint. (**F**) Venn diagram visualizing the number of shared genes between BCG vs. control Day 4 differentially expressed genes in HSC/MPP (p < 0.05), and candidate NRF1-regulated genes.

**Fig. S5.**
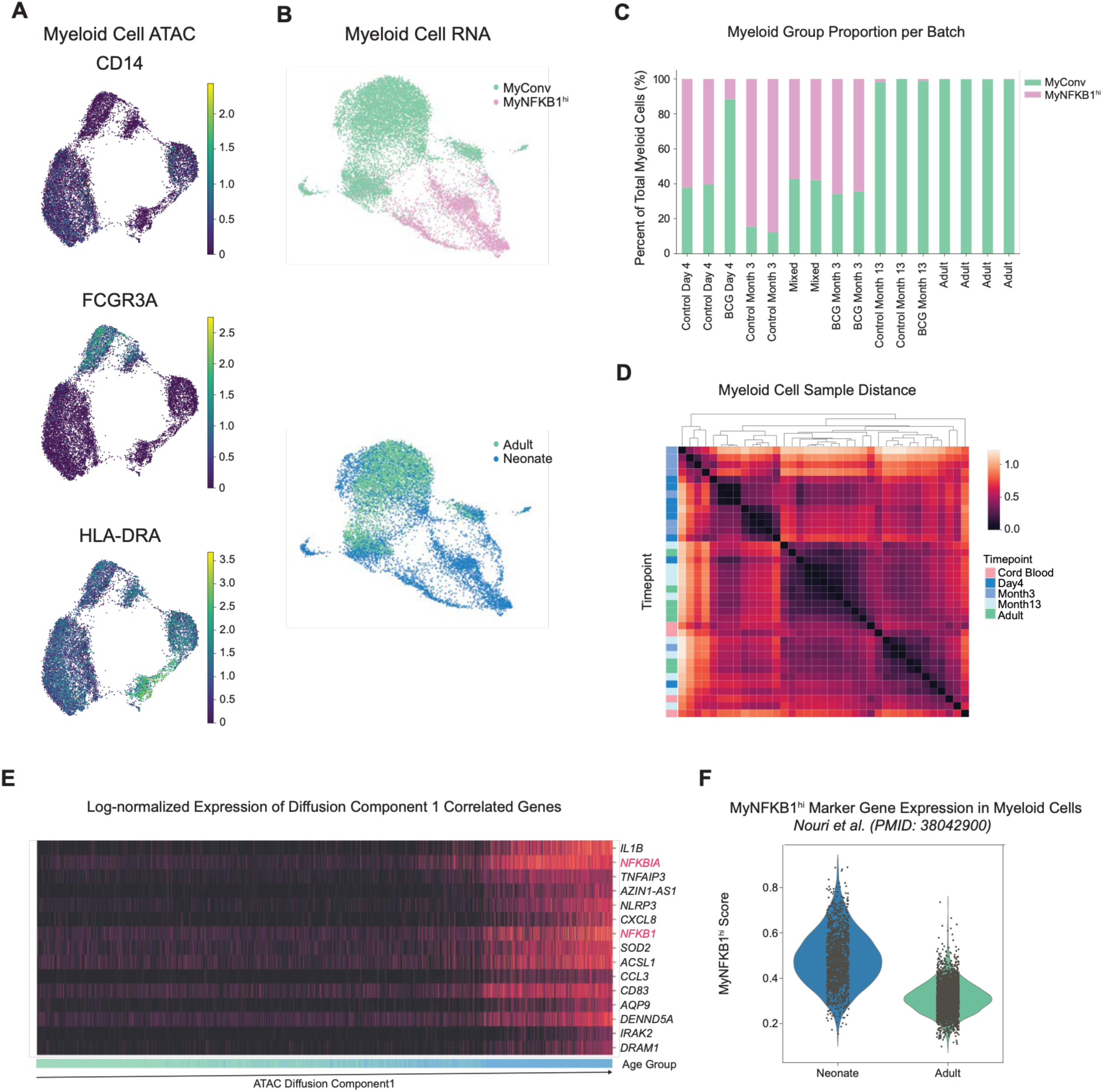
*In-silico* validation of inflammatory myeloid cell signatures in infants. (**A**) UMAP visualization of snATAC data from myeloid cells as in Figure 6B, colored by log-normalized expression of myeloid cell type marker genes. (**B**) UMAP visualization of snRNA data from myeloid cells in all samples, colored by myeloid group annotation (top) and age group (bottom). (**C**) Box plot visualizing proportion of cells annotated to each myeloid group on the y-axis per sequencing batch on the x-axis. (**D**) Heatmap of MrVI sample distance in control myeloid cells. Rows and columns are individual samples, with color value indicating the distance between myeloid cells from each pair of samples in the MrVI *z* latent. Rows are annotated by timepoint. (**E**) Log-normalized expression of Diffusion Component 1 correlated genes (rows) across all myeloid cells (columns) as in Figure 6E. (**F**) Myeloid cells in Nouri et al. 2023 from scRNA sequencing of 2-month old infant PBMC and myeloid cells from 10x scRNA PBMC adult reference data were scored according to expression of MyNFKB1^hi^ marker genes. Violin plot shows distribution of expression scores per age group.

**Fig. S6.**
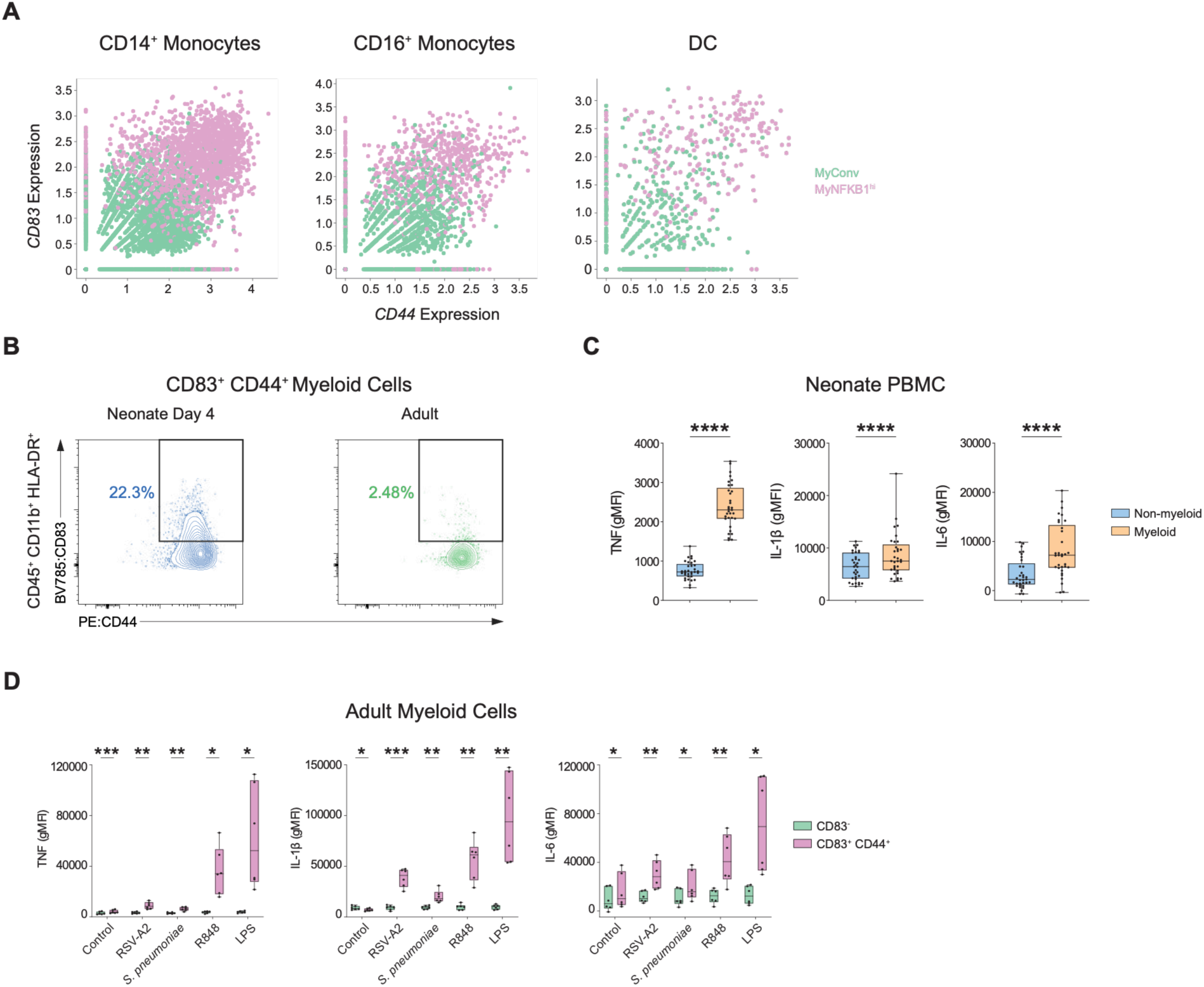
Flow cytometric measurement of CD83^+^CD44^+^ myeloid cells from PBMC. (**A**) Scatter plots of log-normalized snRNA expression of *CD44* vs. expression of *CD83* in PIE-seq data for each myeloid cell type annotation, colored by myeloid group annotation. (**B**) Representative example of flow cytometric analysis of CD83^+^CD44^+^ myeloid subset obtained from neonates at D4 or Adults. (**C**) Box plots depicting geometric mean fluorescence intensity (gMFI) of TNF, IL-1β and IL-6 in neonate myeloid cells vs. non-myeloid cells. *p* values were calculated using a Wilcoxon matched-pairs rank sum test. p > 0.05 = ns, ∗p ≤ 0.05, ∗∗p ≤ 0.01, ∗∗∗p ≤ 0.001, ∗∗∗∗p ≤ 0.0001. (**D**) Box plots of geometric mean fluorescence intensity (gMFI) of TNF, IL-1β, and IL-6 in CD83^+^CD44^+^ myeloid cells vs. non-CD83^+^CD44^+^ after 4h *ex vivo* stimulation with medium control, RSV-A2 (1E05/mL), *S. pneumoniae* (1E08/mL), R848 (10µg/mL), or LPS (10ng/mL) for Adult samples.

**Fig. S7.**
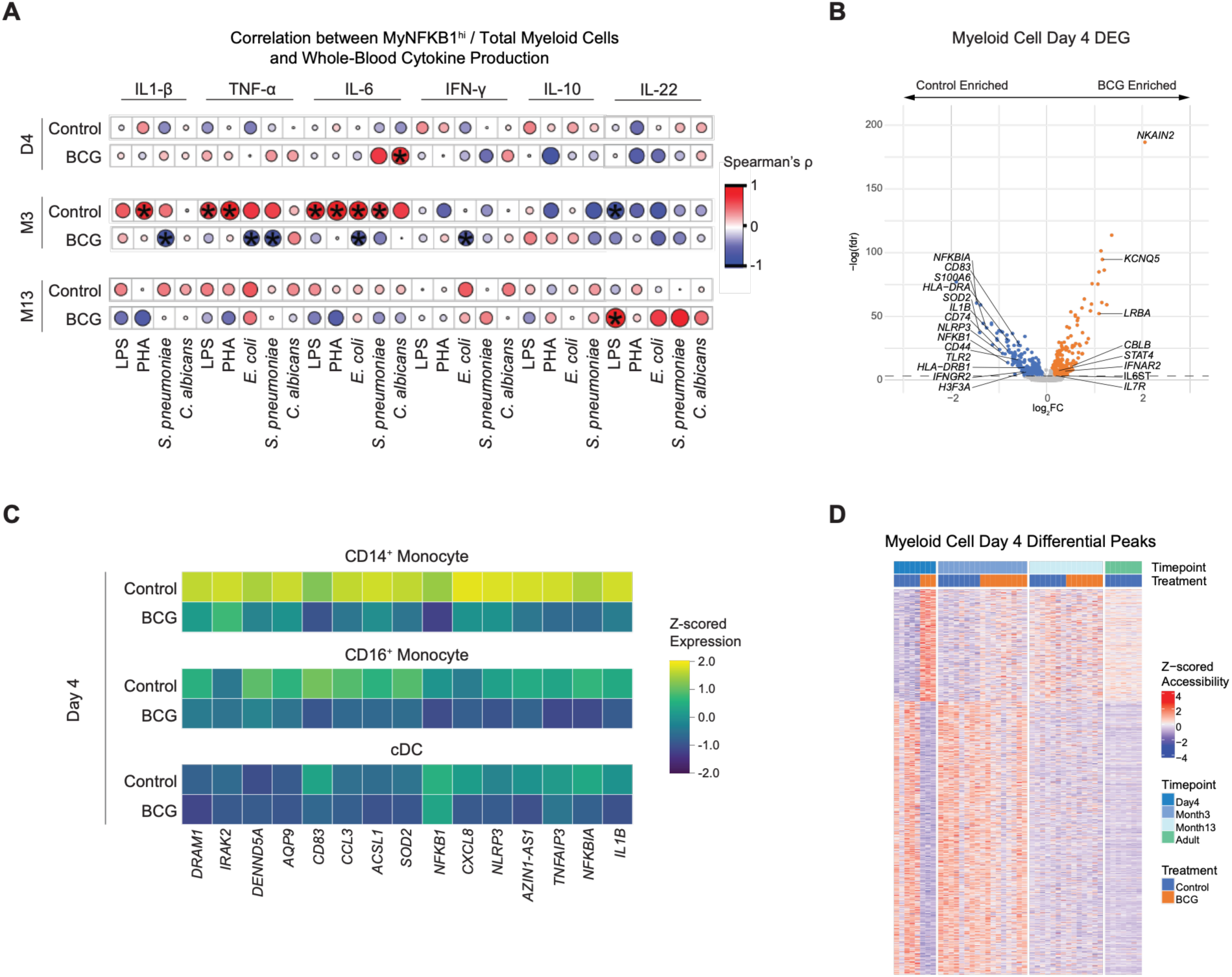
Functional consequences and molecular mechanisms of BCG-associated decrease in neonatal inflammation. (**A**) Dot plot of correlation (Spearman’s ρ) between MyNFKB1^hi^ proportion out of total myeloid cells in the scMultiome dataset and cytokine concentration (pg/mL) in control and BCG matched infant samples 4 days (n = 7 Control, n = 8 BCG), 3 months (n = 8 Control, n = 12 BCG), and 13 months(n = 10 Control, n = 7 BCG) post-randomization upon whole-blood stimulation. Significant (p < 0.05) correlations are marked. (**B**) Differential gene expression results in myeloid cells comparing BCG to control samples at Day 4. Volcano plot shows log_2_FC and -log(FDR) per gene, with genes with abs(log_2_FC) > 0.1 and FDR < 0.001 marked as significant. (**C**) Heatmap of log-normalized gene expression of MyNFKB1^hi^ marker genes (columns) by myeloid cell type and condition (rows) at Day 4. Expression values have been z-scored by column. (**D**) Heatmap of z-scored accessibility at Day 4 BCG vs. control differential peaks (rows) in pseudo-bulk myeloid snATAC data across timepoints and conditions (columns).

**Fig. S8.**
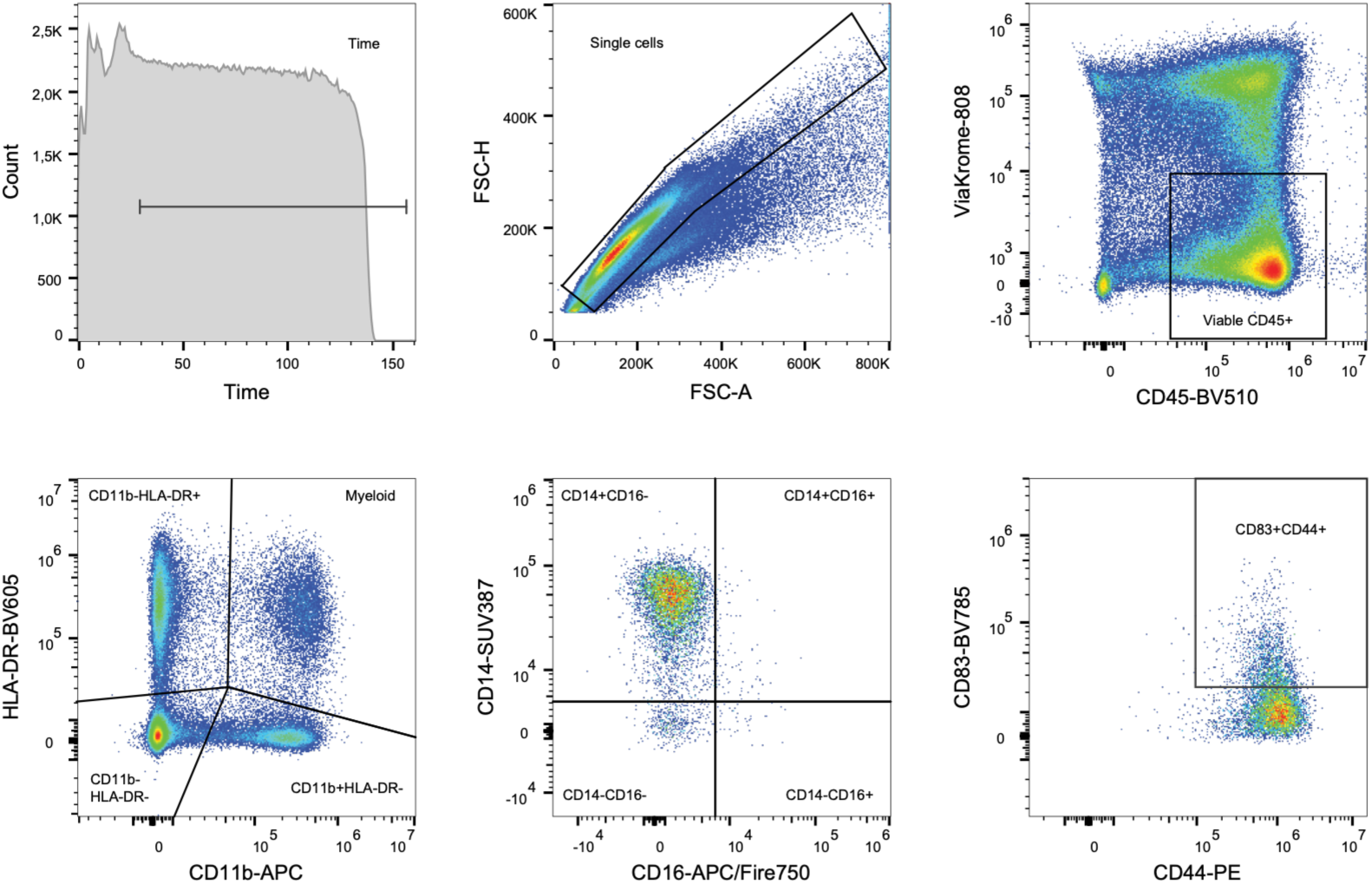
Flow gating strategy for identification of CD83^+^CD44^+^ myeloid cells from PBMC.

